# Assessing SARS-CoV-2 evolution through the analysis of emerging mutations

**DOI:** 10.1101/2022.10.25.513701

**Authors:** Anastasios Mitsigkolas, Nikolaos Pechlivanis, Fotis Psomopoulos

## Abstract

**Intro:** The number of studies on SARS-CoV-2 published on a daily basis is constantly increasing, in an attempt to understand and address the challenges posed by the pandemic in a better way. Most of these studies also include a phylogeny of SARS-CoV-2 as background context, always taking into consideration the latest data in order to construct an updated tree. However, some of these studies have also revealed the difficulties of inferring a reliable phylogeny. [13] have shown that reliable phylogeny is an inherently complex task due to the large number of highly similar sequences, given the relatively low number of mutations evident in each sequence.

**Motivation:** From this viewpoint, there is indeed a challenge and an opportunity in identifying the evolutionary history of the SARS-CoV-2 virus, in order to assist the phylogenetic analysis process as well as support researchers in keeping track of the virus and the course of its characteristic mutations, and in finding patterns of the emerging mutations themselves and the interactions between them. The research question is formulated as follows: Detecting new patterns of co-occurring mutations beyond the strain-specific / strain-defining ones, in SARS-CoV-2 data, through the application of ML methods.

**Aim:** Going beyond the traditional phylogenetic approaches, we will be designing and implementing a clustering method that will effectively create a dendrogram of the involved sequences, based on a feature space defined on the present mutations, rather than the entire sequence. Ultimately, this ML method is tested out in sequences retrieved from public databases and validated using the available metadata as labels. The main goal of the project is to design, implement and evaluate a software that will automatically detect and cluster relevant mutations, that could potentially be used to identify trends in emerging variants.

## 1 Introduction

The greatest combat of the 21st century against a debilitating disease agent SARS-CoV-2 (severe acute respiratory syndrome coronavirus 2) virus discovered in Wuhan, China, in December 2019, has piqued an unprecedented usage of bioinformatics tools in deciphering the molecular characterizations of infectious pathogens. With the viral genome data of SARS-CoV-2 having been made available barely weeks after the reported outbreak, bioinformatics platforms have become an all-time critical tool to earn time in the fight against the disease pandemic.

At the time of writing, there are more than 9.2 million publicly available complete or near-complete genome sequences, that have been submitted to GISAID from all over the world, of severe acute respiratory syndrome coronavirus 2 (SARS-CoV-2) (as of 11 March 2022) and the number continues to grow. This remarkable attainment has been achieved by the inexpensive rapid genome sequencing and online sharing of SARS-CoV-2 genomes by public health experts and research teams around the globe. These genomes play a pivotal role and provide invaluable insights into the ongoing evolution of the virus during the underlying pandemic. Understanding the evolution of not only the SARS-CoV-2 but also other coronaviruses is crucial for creating more robust health infrastructures which prevents major outbreaks under the setting of a pandemic. Although the existence of that “treasure trove” of data, the transition history of the virus and hence “who infected whom” [22] is very difficult to be determined. Previous studies have shown that pinpointing individuals who share the same or almost the same viral genome, could help solve this problem [22] but then there are other difficulties, such as, that we need to identify within-host genetic variants, with high precision, and understand the evolution of them. Needless to say that the phrase “almost the same viral genome” can be poorly defined as there are a lot of ways one can define similarity between two or more sequences and one with physical significance must be preferred.

## 2 Theoretical background

### 2.1 Biological interlude

MicroRNA (abbreviated miRNA) is a small single-stranded non-coding RNA molecule (containing about 22 nucleotides) found in plants, animals, and some viruses, that functions in RNA silencing and post-transcriptional regulation of gene expression [2, 18].

It is widely known that viruses, especially DNA viruses encode miRNAs and, in general, employ the host’s internal machinery of the cells to do so [11]. However, there was a great controversy regarding the existence of miRNAs in RNA viruses, due to the fact that it was theorized that they do not encode miRNAs in order to “avoid excision of their genomes during excision of the precursor miRNAs by the miRNA processing machinery” [24]. Nonetheless, extensive research has focused on that controversy.

Many studies have shown that RNA Viruses, Including CoVs, can encode miRNA-Like small regulatory RNAs. Both [17] and [12] performed deep sequencing using RNA lungs samples from SARS-CoV infected mice and using samples from SARS-CoV and influenza-infected mice strains respectively. They found differentially expressed small ncRNAs—including miRNAs and small viral RNAs (svRNAs) respectively. These virus-derived small RNAs appeared to modulate the host’s response to viral infection by regulating the production of certain pro-inflammatory cytokines (viz. CCL2, interleukin 6, and CXCL10).

The main complication of SARS-CoV-2 infection in a human host, much like SARS-CoV, is the ARDS or acute respiratory distress syndrome, which is caused by an uncontrolled systemic inflammatory response, “cytokine storm” [9], inside the host’s body, and could potentially be deadly.

### 2.2 TSNE

TSNE or t-distributed stochastic neighbor embedding is a stochastic statistical method for reducing the dimensionality of a set of points to low dimensional space [8, 23]. TSNE expresses the similarity and dissimilarity of points in the reduced dimensions by nearby points and very distant points with high probability, respectively.

### 2.3 DBSCAN

DBSCAN or density-based spatial clustering of applications with noise is a density-based non-parametric clustering algorithm utilized in the model building that belongs to the broad group of unsupervised machine learning methods. Given a set of points in an n-dimensional space, DBSCAN is used for grouping together those points which belong to a high-density area, or, in other words, those that have many neighboring points. Points that do not have many neighbors or are pointed out in low-density regions are marked as outliers or noise [7, 20].

### 2.4 MCL

MCL or Markov Cluster Algorithm is an unsupervised clustering method that applies to graphs in the broad concept of what graph means in mathematics. The algorithm makes use of simulation of stochastic flow in graphs. In the field of cluster analysis in graphs, graph partitioning is the process of reducing a graph to smaller graphs by separating its groups of nodes into mutually exclusive groups [3]. The MCL algorithm was invented/discovered by Stijn van Dongen at the Centre for Mathematics and Computer Science (also known as CWI) in the Netherlands [6] and its purpose is to find the optimal partition of a connected graph in an unsupervised way, without any a priori knowledge of the partition sizes.

## 3 Approach

We will approach the problem stated by the research question with the following steps:

- Data gathering
- Data pre-processing
- Data modeling
- Combining binary model with the corresponding metadata to a single file.
- Dimensionality reducing of the binary model for each label at a time. Two different dimensionality reductions (TSNE), one for the non-characteristic (1s) and one for the characteristic (2s) mutations respectively.
- In each one, out of 2, spaces, the one which is constructed by using only 1s and the one which is constructed by using only 2s, we are going to perform unsupervised clustering machine learning, a method called DBSCAN, in an attempt to find high density areas representing closely related samples.
- Up to this point, two completely independent 2-dimensional spaces have been constructed, and on each one, distinct clusters have been identified based on high-density areas. Each cluster represents a group of strongly related samples relying on the characteristic mutations in one space and the non-characteristic mutations in the other space. In other words, we have assigned each sample to a cluster considering only characteristic and non-characteristic mutations each time.
- Undirected graph construction based on co-occurring mutations over groups of samples as they were indicated based on the clusters of the previous step.
- Finding highly interconnected parts of the network through MCL analysis.
- Directed graph construction based on occurring mutations from one node to others. This graph should indicate evolutionary paths of the virus.
- Validating some possible evolutionary paths by using the MSA of those samples that take part of those paths and finding correlations among their co-occurring mutations.
- Using these correlations as a measure of distance and run hierarchical clustering.
- The result dendrogram should indicate strongly related co-occurring mutations among a group of samples involving an evolutionary path.

## 4 Materials and methods

In this study, we present a novel method for detecting patterns of co-occuring mutations beyond strain-specific / strain-defining ones and making use of those patterns in an endeavor to group samples in such a way that they could indicate evolutionary paths of the virus. In otherwords, mutational occurrence patterns might suggest different ways of grouping sample revealing the evolutionary history of SARS-CoV-2.

### 4.1 Data collection

For this study, 5411 vcf files and their metadata have been deposited in the European Nucleotide Archive (ENA) at EMBL-EBI under accession number PRJEB44141 ENA. Severe acute respiratory syndrome coronavirus 2 isolate Wuhan-Hu-1, complete genome with NCBI Reference Sequence: NC_045512.2 was obtained from NCBI database and used for the alignment of the raw data, the variant calling, the consensus sequence generation (alternative genomes), and the remaining algorithmic procedures described in the rest of this study. At this point, it is worth mentioning that the raw sequences alignment and the variant calling were not part of this study and had been performed previously. For the time being, we obtained the output (vcf files [5]) of the variant calling pipeline. A Genetic variant annotation and functional effect prediction toolbox, SnpEff [4], was then used in order to annotate all the vcf files, at the amino acid level.

### 4.2 Data summary

The dataset consists of 5411 vcf files and their metadata. Each file represents a sample collected by the health care system in Greece and altogether covers a period of the second COVID-19 pandemic wave, from 15-04-2021 to 15-08-2021 (figure 3). In order to eliminate the sampling bias and the sequencing errors, duplicate and high ambiguity sequences were removed. To achieve that, the pangolin tool was employed and only the sequences able to be classified were kept. A summary of the assigned lineages to the used sequences is given in table 1. The kept 4.500 sequences were then aligned to the SARS-CoV-2 genome using MAFT [10] and were used for the rest of the algorithmic procedures, as explained in detail in the next chapters.

**Fig. 1.**
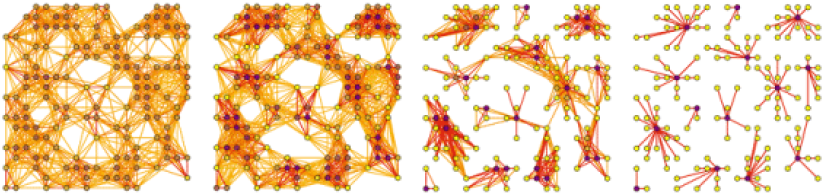
Markov clustering principal illustration. Copied from: http://micans.org/mcl

**Fig. 2.**
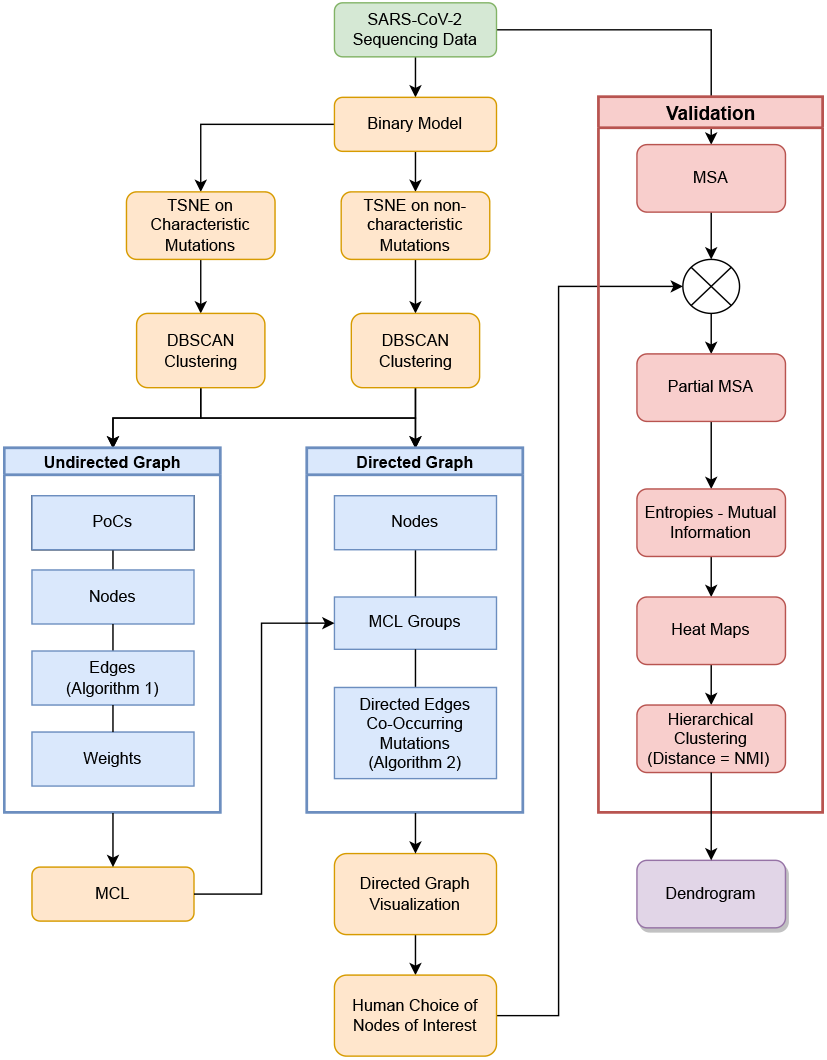
Flow diagram. Materials and methods flow illustration.

**Fig. 3.**
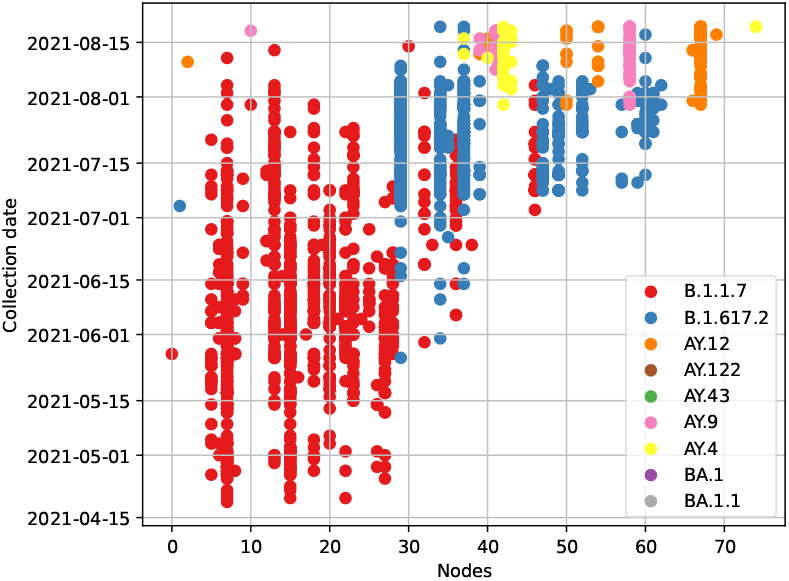
Samples ordered by collection date. Each dot in this graph represents a sample and its corresponding collection date is shown by the y-axis. The node that a particular sample belongs to is shown by the x-axis and the assigned lineages by the pangolin classifier are shown in different colors.

**Table 1.** Dataset summary

| Pango Lineage | WHO Label | Description | Occurrences |
| --- | --- | --- | --- |
| B.1.1.7 | $\alpha$ Alpha | UK lineage of concern | 1278 |
| B.1.617.2 | $\delta$ Delta | Predominantly India lineage | 1018 |
| AY.43 |  | Alias of B.1.617.2.43, European lineage | 512 |
| BA.1 | $\omicron$ Omicron | Alias of B.1.1.529.1, from pango | 431 |
| AY.12 |  | Reassigned to AY.122.1 | 285 |
| AY.4 |  | Alias of B.1.617.2.4, UK lineage | 228 |
| AY.122 |  | Alias of B.1.617.2.122, European lineage | 221 |
| AY.9 |  | Alias of B.1.617.2.9, UK lineage | 120 |
| BA.1.1 |  | Alias of B.1.1.529.1.1 | 96 |
| AY.125 |  | Alias of B.1.617.2.125, European lineage | 59 |
| AY.9.2 |  | Alias of B.1.617.2.9.2, European lineage | 53 |
| B.1.1.318 |  | Lineage circulating in multiple countries with a number of variants of concern (Spike D614G, Spike D796H, Spike E484K, Spike P681H, Spike T95I, Spike Y144del). | 52 |
| B.1.1.529 | $\omicron$ Omicron | South Africa and Botswana lineage | 46 |
| AY.121 |  | Alias of B.1.617.2.121, European lineage | 45 |
| AY.4.1 |  | Alias of B.1.617.2.4.1, UK lineage | 30 |
| AY.120 |  | Alias of B.1.617.2.120, UK lineage | 28 |
| AY.42 |  | Alias of B.1.617.2.42, European lineage | 27 |
| AY.7 |  | Alias of B.1.617.2.7, UK lineage | 25 |
| AY.84 |  | Alias of B.1.617.2.84, European lineage | 12 |
| AY.39 |  | Alias of B.1.617.2.39, predom. USA lineage | 10 |
| AY.126 |  | Alias of B.1.617.2.126, European lineage | 10 |

This table shows a summary of the samples used in this study. Starting from the left, the first column shows the Pango Lineage name [19]. The second one shows the WHO Labels while the third one gives a description, mostly, regarding the area of dominance of those lineages. The fourth column is the number of each lineage appearing in the entire dataset [14].

A vast amount of invaluable information, such as the sample id, collection date, geographic location and/or geographic region, taxonomy, host sex, a small description of how the viral sequences were retrieved from human hosts, and other information is provided in the form of metadata in XML format by ENA database. A script in python was then used to clean and transform the XMF format to an easy-to-read in human-readable CSV format. In addition to those fields, the dynamic nomenclature proposal for SARS-CoV-2, given by the pangolin tool is provided for each sample, as a completely separate CSV file. Since we merged those CSVs for simplicity reasons, a detailed list with the exact names of columns in the CSV metadata file is provided in Table 2, which must be followed exactly by the reader in case one would like to reproduce the results.

**Table 2.** Metadata CSV file column names

| Column name | Description |
| --- | --- |
| sample name | eg. Vxxx (xxx is an integer number) |
| collection date | The collection date of the sample |
| geographic location | The country in which the sampling took place |
| geographic location | Region and locality (same as above) |
| host common name | eg. Human |
| host scientific name | eg. Homo sapiens |
| host sex | male/female/not provided |
| host health state | If provided |
| isolate | How was it isolated |
| lineage | The dynamic nomenclature for SARS-CoV-2 lineages by pangolin tool |

This table shows the metadata CSV file column names. They must be strictly followed as such, in order for one to be able to reproduce the results.

### 4.3 Modeling

A data model is an abstract mathematical representation that assembles the elements of a dataset in terms of a standard or a template format. Data models are also used to describe the data themselves, how the elements are related to one another, and the properties of a real-world problem.

As we have already described in the above sections, our dataset consists of raw SARS-CoV-2 genomes. We can have access to these raw sequences either by looking at the FASTA file which contains the actual sequences or by accessing the VCF files where only the specific mutations and their characteristics are stated, in correspondence with the reference genome. The reason why we need a model to represent our data is becoming more and more clear. Given that the data is unstructured and quite complex to be analyzed by different algorithms we needed a very simple model or a mathematical representation of them, very well structured and easy to manipulate. Preferably a model that consumes a small amount of computer memory, not only because we have a few thousand genomes but also because our initial goal was to provide a tool as generic as possible, capable of analyzing large datasets on a personal computer or even a laptop.

At this point, we will be explaining the model that we chose to work with, its advantages and disadvantages over other models that exist in the bibliography, and most importantly what are the necessary assumptions one has to take for granted to make use of such a model.

Given the fact that the mutations’ locations are given relative to the reference genome, one could model the raw – mutated sequences using a binary model of the same length as that of the reference genome. Simply put, one shall use a string representation, per sequence, with the same length as that of the reference genome. This string must be capable of encoding all the different possible states that a site can physically exist in. In our case three different symbols are being used, zeros, ones, and twos, one for each of the three different states, unmutated sites, characteristic mutations as reported by the pangolin tool, and non-characteristic mutations Fig. 4. Although the abovementioned model is not exactly binary, because of the use of three symbols instead of two, from now on we are going to call it binary as one can use a combination of only the first two symbols, zeros, and ones, to describe the third state. The decision of using three symbols was made by us just to reassure a crystal-clear distinction among the states. It is worth mentioning that by using such a model and by keeping the length of the string representations constant relative to the ref genome, only the deletions and the substitutions can be taken into consideration while the insertions must be discarded. Otherwise, the length of the model will not be the same for each sequence. As a result, our binary model lacks the ability of modeling insertions, and this is indeed its main drawback. Despite the loss of some information, this type of modeling method seems sufficient enough, at least at the time of writing, since a careful examination of the characteristic mutations reported by the pangolin tool, showed that not too many insertions had been reported. Hence, we can safely use a model that is only able to model deletions and substitutions.

**Fig. 4.**
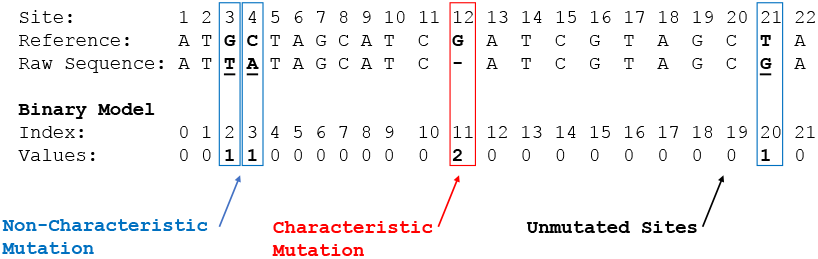
Example illustrating how the bin model is used. Given a raw sequence, mutated sites are represented with “1” or “2” according to whether they are non-characteristic or characteristic mutations, respectively. Unmutated sites are represented with “0”.

### 4.4 Dimensionality reduction and clustering on the binary model

Due to the nature of the problem of modeling and analyzing raw sequences represented by a binary model, there are many ways to manipulate the data regarding either the columns of the binary table as features and the rows as records or vice versa. In an attempt to identify correlations among different coordinates or simply put sites, of the virus genome, at least completely intuitively, one could consider defining each site of the genome as a variable and the samples (rows of the table) as records. We end up with a dataset that consists of almost 30k variables, as many as the sites are, which without any further modifications is quite difficult for someone to manipulate because of its high dimensionality. On the other hand, one could consider each site as a variable that changes as we move along the samples. In other words, each variable could be considered as a time series that changes while we are moving from one sample to the next, under the assumption that the samples are in sequential temporal order Fig. 6A, which as we have already discussed is impossible for one to know in a pandemic setting.

**Fig. 5.**
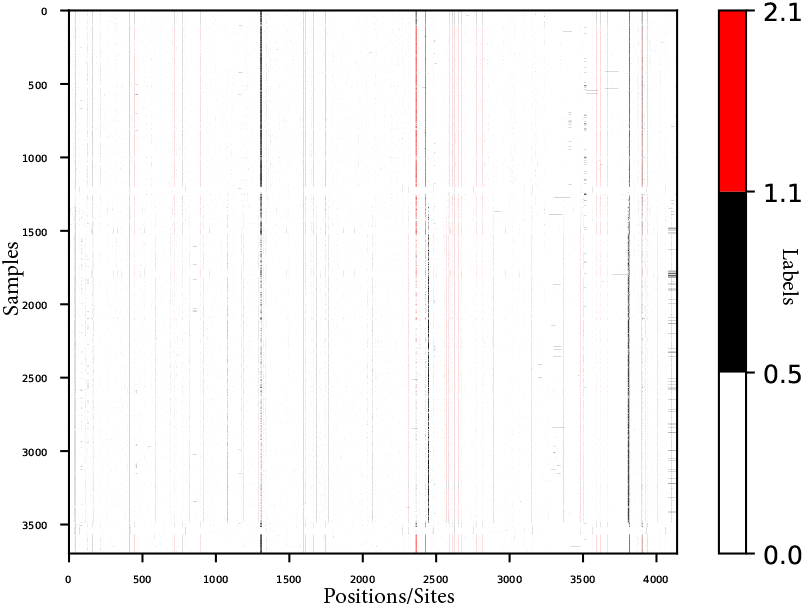
Binary model along ~ 4000 different samples. The y-axis represents the sample indices while the mutated sites are shown in the x-axis. Note that the x-axis does not show the actual site positions but a simple counter. While the SARS-CoV-2 genome length is ~ 30*k* sites, only the 4500 of them that appear to be mutated at least once within the dataset, are shown in this graph. The characteristic pangolin-based mutations are shown in red while the non-characteristic ones are shown in black pixels. The rest, un-mutated sites appear as white pixels.

**Fig. 6.**
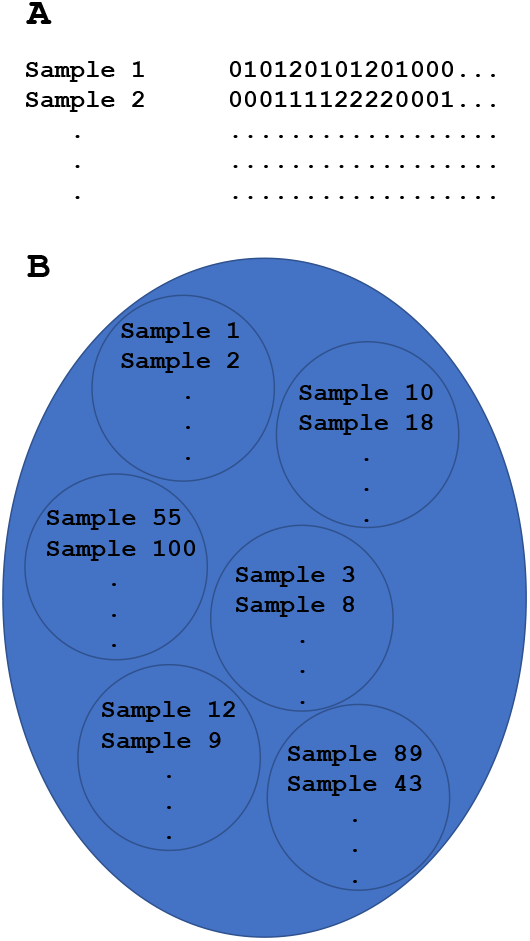
(A) Samples in sequencial temporal order. (B) Samples clustering. Each cluster represents closely related samples.

In order to visualize and make the dataset more understandable, a variety of dimensionality reduction methods were performed. A comparison of them is not going to be mentioned since that would be beyond the scope of this report. TSNE with two components was employed for the task of lowering the dimensions of the data, providing a 2-dimensional representation, and making it easier for us to visualize, understand, and form clusters of closely related samples. Besides TSNE performed best in comparison with other methods, in both the ability to separate distant samples and the execution time.

In an attempt to produce two completely different and independent 2D spaces, one able to separate samples based on the characteristic mutations and one based on the non-characteristic mutations, TSNE was used twice, one taking into account only the non-characteristic mutations and one taking into account only the characteristic mutations. As a result, two different 2D spaces have been calculated, the first based on the characters “1” Fig. 7 and the second based on the characters “2” Fig. 8. From now on, we will be using the notation “based on 1s” or “based on 2s” when we want to refer to the clustering made by considering only the non-characteristic or characteristic mutations respectively.

**Fig. 7.**
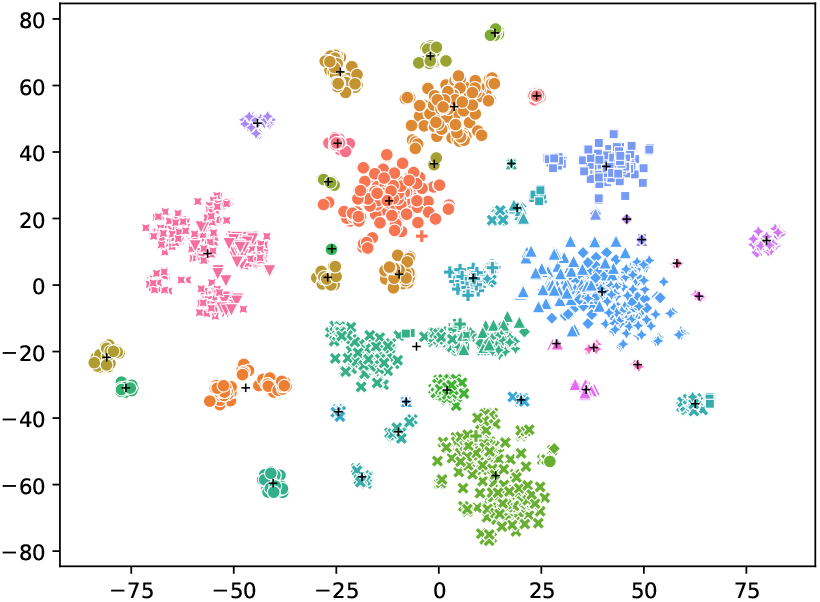
DBSCAN on TSNE 2D dimensional space, considering only non-characteristic mutations. Individual axes in this graph have no meaning at all and that is why their labels are not shown. Different lineages are shown with different symbols. As a result of DBSCAN, each cluster is shown in a different color while the clusters’ centroids are shown by black crosses.

**Fig. 8.**
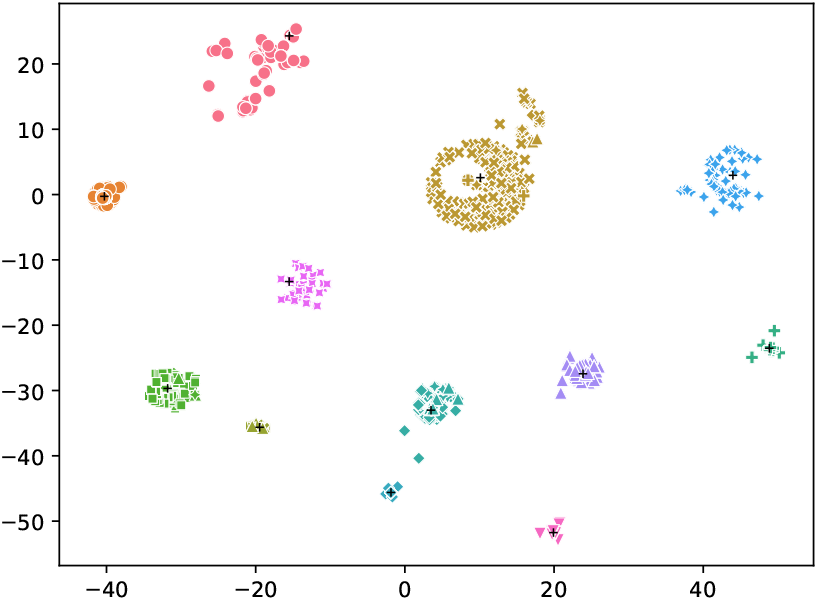
DBSCAN on TSNE 2D dimensional space, considering only characteristic mutations. Individual axes in this graph have no meaning at all and that is why their labels are not shown. Different lineages are shown with different symbols. As a result of DBSCAN, each cluster is shown in a different color while the clusters’ centroids are shown by black crosses.

Regarding the 2D space of the data considering only the characteristic mutations Fig. 8, one can clearly distinguish how well separated the clusters of different lineages are. The fact that the groups of lineages are almost perfectly separated from each other is profound but not completely unexpected. This is proof of not only how powerful TSNE could be in separating sequences in some cases but also how precisely the mutations are selected as the main features of the pangolin classifier since they separate the sequences in groups in such a great way.

Since TSNE keeps closely related samples nearby, within the 2D space, the next logical step is to use a necessarily unsupervised machine learning technique to identify clusters of interest in the reduced dimensional spaces. It is of great necessity to build up an automated method of identifying high-density clusters, even if those clusters consist entirely of samples that have been classified as they belong to the same lineages. The reason why, as it has already been mentioned, is that it is very challenging to put the samples in sequential temporal order and, therefore, groups of closely related samples must be formed assuming that a “jump” between nearby groups can indicate a possible underlying evolutionary path of the virus.

For that purpose, Scikit Learn implementation of the DBSCAN algorithm was used with the default interface parameters except for the “eps” value. EPS is defined as “the maximum distance between two samples for one to be considered in the neighborhood of the other” [16] and set up equal to 5 using a simple trial and error approach. DBSCAN performed almost perfectly in the case of characteristic mutations and satisfactorily in the case of non-characteristic mutations as shown in Fig. 7, since most of the large distinct groups of the same lineage have been identified and high-density sub-areas within groups of the same lineage have been caught.

### 4.5 Graph construction

#### Positions of concern (PoCs)

As a result, each cluster, regardless of whether they were formed based on “1”s or “2”s, consists of a bunch of samples and their assigned lineages (see section 4.4).

We define a PoC as the position of a nucleotide that appears to be mutated with a non-zero frequency, along the samples of the same lineage of a very particular cluster based on “1”s or “2”s. As a result, we construct a table, the column names of which follow the bellow mentioned notation:

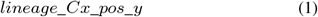

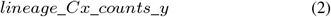

where:

- *lineage* is the lienage;
- *x* is the unique identifier of the particular cluster;
- *y* is an index that shows based on which symbol the clustering has been done.

The columns named by equation 1 provide the mutated sites while the columns named by equation 2 provide the counts for those sites. Once again, only the sites with non-zero counts or *frequency* > 0 are being stored in this table.

##### Example 4.1.

*Let’s assume lineage = B*.1.1.7, *x* =1, *and y* =2, *which means that we are looking for the PoCs of samples with lineage B.1.1.7 that belong to the cluster 1 based on “2”s. Given the equations 1, 2, the two column names will become*:

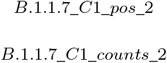

*and thus, the PoCs table will be:*

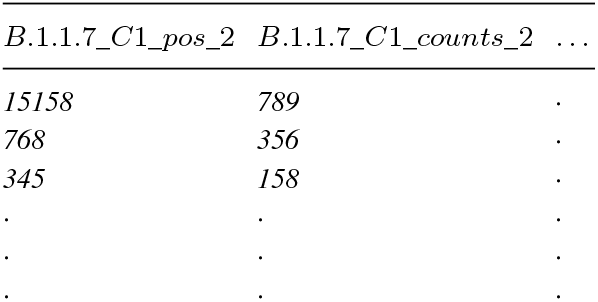

| $B.1.1.7\_C1\_pos\_2$ | $B.1.1.7\_C1\_counts\_2$ | ... |
| --- | --- | --- |
| 15158 | 789 | . |
| 768 | 356 | . |
| 345 | 158 | . |
| . | . | . |
| . | . | . |
| . | . | . |

***Each data mentioned in this example is just dummy values and does not represent any real information.***

#### Nodes

Let

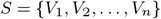

the collection of all samples,

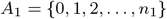

the collection of the unique clusters’ identifiers based on 1s (noncharacteristic mutations),

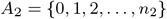

the collection of the unique clusters’ identifiers based on 2s (characteristic mutations),

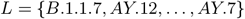

the collection of possible lineages in the dataset, with

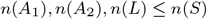

where *n*(*X*) is the number of elements in set *X*. Let also *M*_1_ : *S* → *A*_1_, *M*_2_ : *S* → *A*_2_ and *M*_3_ : *S* → *L* be three maps, that given a particular *s_i_* return the cluster in which the sample belongs to, based on “1”s and “2”s, and its assigned lineage, respectively. Table 3 shows the way the abovementioned sets are structured. We define, for each (*a*_1_, *a*_2_) ∈ *A*_1_ × *A*_2_, the following subset of *S*:

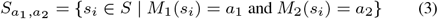

and the following subset of *L*:

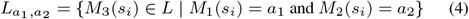

**Table 3.** Samples

| sample $s_i$ | cluster_1 $M_1(s_i)$ | cluster_2 $M_2(s_i)$ | lineage $M_3(s_i)$ |
| --- | --- | --- | --- |
| $V_1$ | 0 | 2 | AY.12 |
| $V_2$ | 1 | 2 | B.1.1.7 |
| $V_3$ | 3 | 0 | AY.7 |
| $V_4$ | 1 | 2 | AY.12 |
| $V_5$ | 1 | 2 | B.1.1.7 |
| . | . | . | . |
| . | . | . | . |
| . | . | . | . |

The *S*_*a*_1_, *a*_2__, as defined by equation 3, is a subset of *S*, whose elements are all the samples that have been assigned to these particular *a*_1_, *a*_2_ clusters, while the *L*_*a*_1_, *a*_2__, as defined by equation 4, is a subset of *L*, whose elements are all the assigned lineages of the corresponding samples *S*_*a*_1_, *a*_2__.

Given each combination of clusters (*a*_1_, *a*_2_), and the lineages *L*_*a*_1_, *a*_2__ of the included samples, one can easily use the *a*_1_, *a*_2_, *L*_*a*_1_, *a*_2__, the equations 1, 2, and the PoCs table in order to determine the *PoCs*_*a*_1__, *PoCs*_*a*_2__ for each *lineage* ∈ *L*_*a*_1_, *a*_2__. After that, we used for each *lineage* ∈ *L*_*a*_1_, *a*_2__, the PoCs of *a*_1_ and *a*_2_, separately, (clarification: saying *a*_1_ and *a*_2_, means, cluster *a*_1_ based on “1”s and cluster *a*_2_ based on “2”s) in an attempt to find out which of these positions of concern would appear as SNPs in the samples *S*_*a*_1_, *a*_2__, by searching for SNPs at the *PoCs*_*a*_1__, *PoCs*_*a*_2__ along the samples *S*_*a*_1_, *a*_2__. When a SNP, the position of which ∈ *PoCs*_*a*_1__ or *PoCs*_*a*_2__, is found among the samples *S*_*a*_1_, *a*_2__, a new record is added to the data structure (table 4) to the -based on “1”s snps- or -based on “2”s snps-column, regarding whether the position of that SNP belongs to the *PoCs*_*a*_1__ or *PoCs*_*a*_2__.

**Table 4.**
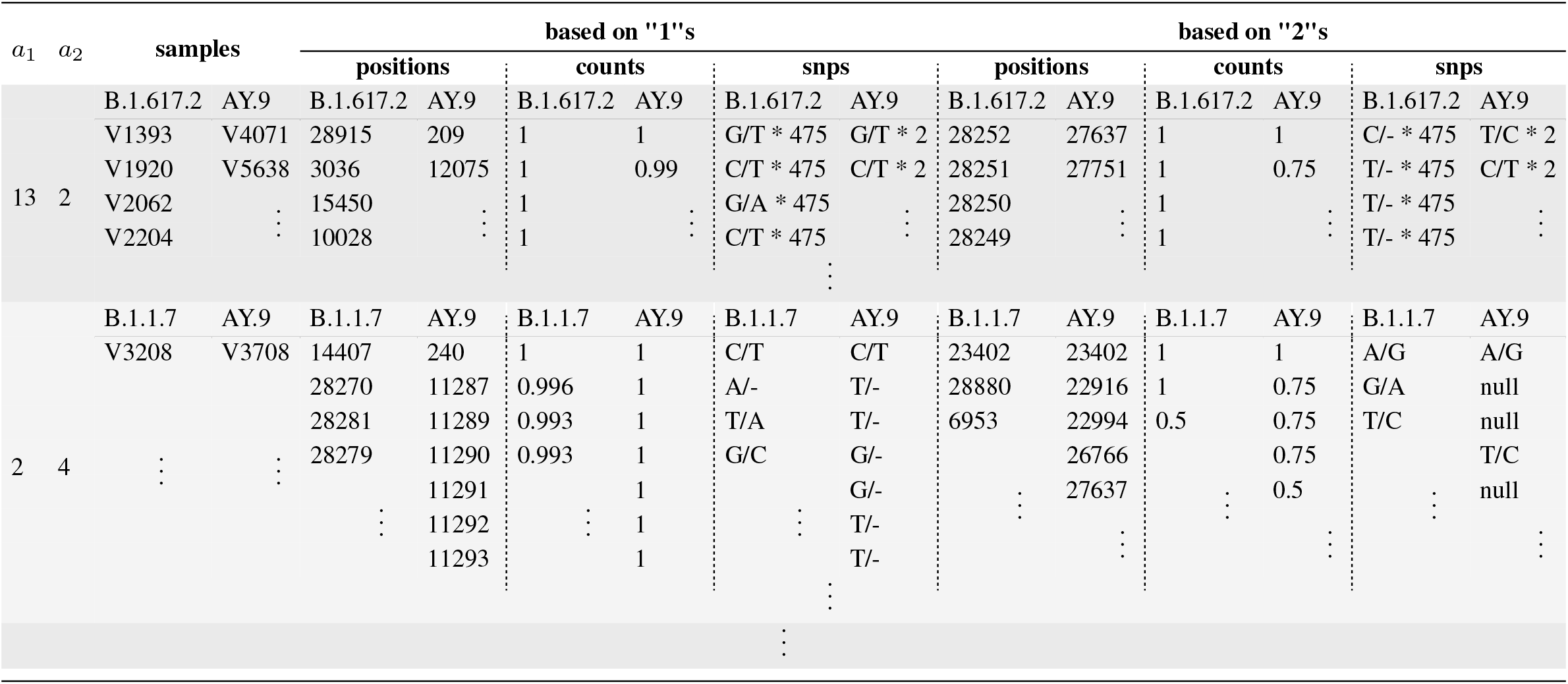
Example data. This table illustrates the data structure used to store the complex objects of nodes. In this table, each row (different shades of gray) represents a node.

#### Edges

We define:

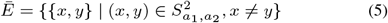

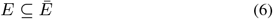

While the cartesian product of *S*_*a*_1_, *a*_2__ with itself, denoted 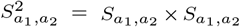 is the set of all ordered pairs (*x, y*), the set 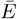 is defined above, equation 5, as the cartesian product of *S*_*a*_1_, *a*_2__ with itself using only unordered pairs under the constraint *x* ≠ *y. E* is simply a subset of 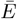, the purpose of which will become clear later in this chapter.

##### Example 4.2.

*Supposing S*_*a*_1_, *a*_2__ = {*a,b*}, *the cartesian product of S*_*a*_1_, *a*_2__ *with itself will be:*

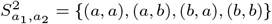

*According to the equation 5*, 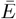, *which once again is the cartesian product of S*_*a*_1_, *a*_2__ *with itself using only unordered pairs under the constraint x* ≠ *y, will be*:

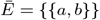

*and simply:*

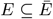

*a subset of* 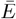.

In that way, we have constructed set 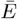, whose elements are the combinations of elements of *S*_*a*_1_, *a*_2__, as unordered pairs without including pairs of the same elements. To put it simply, given that *S*_*a*_1_, *a*_2__ is composed by the records of the table 4, 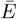 will give us all the unique combinations among them, in pairs of two, without taking into consideration the order and the pairs of an element with itself.

As we will see in detail in the following section, assuming that the elements of *S*_*a*_1_, *a*_2__ are nodes in an undirected simple graph, then *E* is a set of edges, each of which is associated with two distinct nodes.

Now, let’s come back to the definition of edges. Even though set 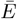 has already been expressed in a set-builder notation and we have a well-defined mathematical notation of this set to work with, we haven’t mentioned yet, how the elements of the subset *E* are supposed to be selected from set 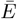 of equation 5. For that purpose algorithm 1 is used.

Given the combinations of nodes, 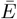, as defined by equation 5, algorithm 1, for each pair of nodes 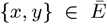 (for loop in line 1), retrieves lineages *L_x_, L_y_* (line 3,4)and for each lineage *l_x_* ∈ *L_x_*, *l_y_* ∈ *L_y_* calculates the common positions of concern (for loops in lines 5,6). Once these loops over lineages *L_x_* and *L_y_* are done the algorithm is responsible for checking whether or not there has been at least one position of concern in common (line 12). If indeed one or more positions of concern is found in common within those nodes {*x, y*} then an edge between those two is being considered and stored in subset *E* (line 13). The weight of that edge is defined as the number of the common positions of concern (line 14). It now becomes clear why *E* is a subset of 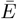. That is because the algorithm connects with edges only those pairs of nodes {*x, y*} in which there is at least one common position of concern.

Miscellaneous calculations are not shown in algorithm 1 for simplicity reasons. In the real implementation of that algorithm many data, beyond the weight, accompanying each edge {*x, y*} are calculated as well. For instance, a small table accompanies each edge, containing for each given *l_x_, l_y_* pair the common *PoCs_x, y_* of the connected nodes, and their minimum occurrence frequency.

##### Algorithm 1 Calculate undirected edges

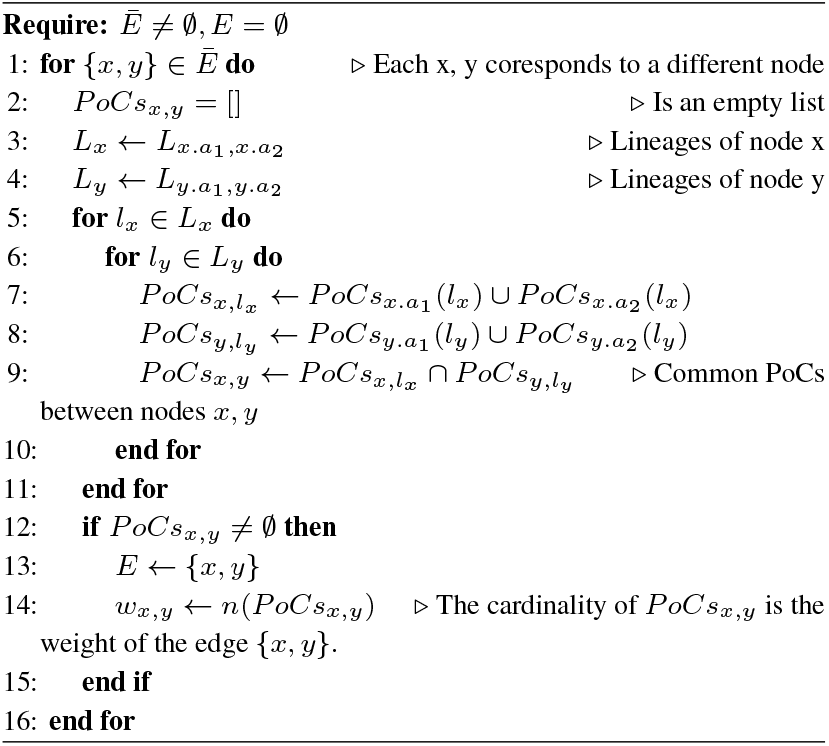

##### Example 4.3.

*Given an element* {*x, y*} *of E, which represents an edge connecting nodes x and y, the following PoCs_x, y_ have been found in common:*

| based on "1"s |  |  |  | based on "2"s |  |  |  |
| --- | --- | --- | --- | --- | --- | --- | --- |
| positions |  | counts |  | positions |  | counts |  |
| AY.7 | AY.9 | AY.7 | AY.9 | AY.7 | AY.9 | AY.7 | AY.9 |
| 28915 | 209 | 1 | 1 | 28252 | 27637 | 1 | 1 |
| 15450 |  | 1 |  | 28250 |  | 1 |  |
| 10028 |  | 1 |  | 10028 |  | 0.5 |  |
| B.1.1.7 | AY.9 | B.1.1.7 | AY.9 | B.1.1.7 | AY.9 | B.1.1.7 | AY.9 |
| 14407 | 240 | 1 | 1 | 23402 | 23402 | 1 | 1 |
| 28270 | 11287 | 0.996 | 1 | 28880 | 22916 | 1 | 0.75 |
| 28281 | 11289 | 0.993 | 1 | 6953 | 22994 | 0.5 | 0.75 |
| 28279 | 209 | 0.993 | 0.75 | 10028 | 26766 | 0.4 | 0.75 |
|  | 11291 |  | 1 |  | 209 |  | 0.5 |
|  | 11292 |  | 1 |  |  |  |  |

*Where:*

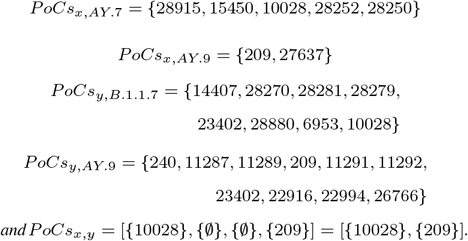

*This means that the edge which connects nodes* {*x, y*} *will have got weight: w_x, y_* = *n*(*PoCs_x, y_*) = 2 *and the accompanying table will be:*

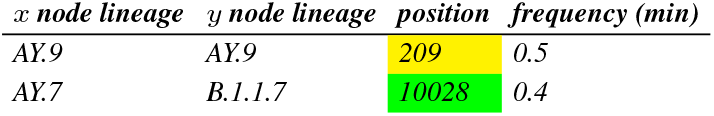

***Each data mentioned in this example is just dummy values and does not represent any real information.***

##### Graph definition

In the previous chapters, *S*_*a*_1_, *a*_2__ was defined, which for a given tuple (*a*_1_, *a*_2_) represents a node of a graph. Each of those nodes is composed by a number of samples of some given lineages. These samples belong to the corresponding tuple of clusters (*a*_1_, *a*_2_) based on “1”s, and “2”s, respectively. In addition, we have defined *E*, a set whose elements are carefully selected edges, each of which connects two distinct nodes, as described in the previous chapter. The edges *E* have been selected in such a way, that they connect a pair of two nodes if and only if those nodes are composed by at least one position of concern in common.

Let

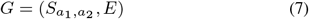

be an undirected graph, or in other words a tuple comprising:

- *S*_*a*_1_, *a*_2__, a set whose elements are called vertices (or nodes);
- *E*, a set of paired vertices, whose elements are called edges.

Despite the fact that the *S*_*a*_1_, *a*_2__ and the *E* are used in order to define the graph (see eq. 7) and, by definition, these two sets consist of a group of samples for each element of *S*_*a*_1_, *a*_2__ and an edge connecting two elements of *S*_*a*_1_, *a*_2__ for each element of *E*, in reality, as has already been mentioned, both the nodes and the edges are accompanied by additional supplementary information.

More specifically, with regards to the nodes, for each one of them:

- the samples *S*_*a*_1_, *a*_2__ of which the node is made up;
- the lineages *L*_*a*_1_, *a*_2__ assigned to those samples;
- the percentage content of the compound lineages (E.g.: If in a particular node 50 out of 100 samples are B.1.1.7 and the rest 50 are AY.7, the percentage content will be 50-50%.);
- the positions of concern *PoCs*_*a*_1__ and *PoCs*_*a*_2__ for each of the assigned lineages;
- the frequencies or counts of the positions of concern, as they appear from the point of view from the inside of cluster *a*_1_ and *a*_2_ (not from inside the samples of this particular node);
- the SNPs that were found at either some or all the positions of concern, as well as how many in each position;

are calculated and shown in a small table that accompanies a corresponding node.

As for the edges, they are accompanied by a corresponding table such as that already mentioned in example 4.3. The connecting power of an edge is given, by definition, by its weight and it is shown as the thickness of the line which connects two distinct nodes {*x, y*}, for visualization purposes.

A very convenient graph UI might be used, so that, when a user hovers the mouse over a node/edge all the necessary information appears in a pop-up table. To represent the constructed graph in a better way, Pyvis, a python library for visualizing networks, is used. That library is capable of using small icons instead of regular circles as nodes and variable thickness for the edges connecting nodes between them, which makes it extremely useful for our purposes.

### 4.6 MCL

By visually examining the structure of the produced network, we end up with a few rather obvious conclusions. First, the graph seems to be almost fully connected. That should be explained if one considers that in our thoroughly explained dataset, the mutation rate is relatively low compared with the number of samples, and the majority of the non-characteristic mutations is circulating among them. This brings us to the conclusion that between two groups of different samples (graph’s nodes) there should be at least some mutations in common and thus we end up with an edge connecting those two nodes. As seems to be the case, the high interconnectivity of nodes leads to a highly interconnected graph. Second, and due to the fact that the graph appears to be fully connected, it is very intricate to draw conclusions and even harder for one to intuitively recognize high-density clusters of interconnected nodes.

For the above-mentioned reasons, there is a great necessity of using a computational method to formulate clusters of high-density groups of nodes in that graph. MCL is mobilized to do so. We ran the MCL algorithm using the default input parameters on our model graph, in an attempt to identify clusters and prune some edges among them. The ultimate goal was to end up with a rather not fully connected network. One could say that the aim is to “relax” strong bonds among these groups of nodes ending up with strongly interconnected groups of nodes, weakly connected to each other.

MCL takes into consideration graph *G* as defined by equation 7 and the weights *w_x, y_* of each pair of nodes {*x, y*}. According to the documentation, MCL is unequivocally able to process weighted undirected graphs and the weights must be non-negative numbers while it is explicitly stated that they have to represent similarities instead of dissimilarities. With that being said and since our weights, as defined in the previous chapter, are positive integers, MCL took place. In the first few attempts, the results were unexpectedly bad since the MCL was incapable of distinguishing any clusters at all. It was soon made clear that the underlying problem lay in how the weights were distributed rather than the MCL itself.

Since an edge’s weight represents the number of common mutations for each pair of lineages, between the nodes {*x, y*}, one could consider the *w_x, y_* as a variable, the distribution of which is given by Fig. 10.

**Fig. 9.**
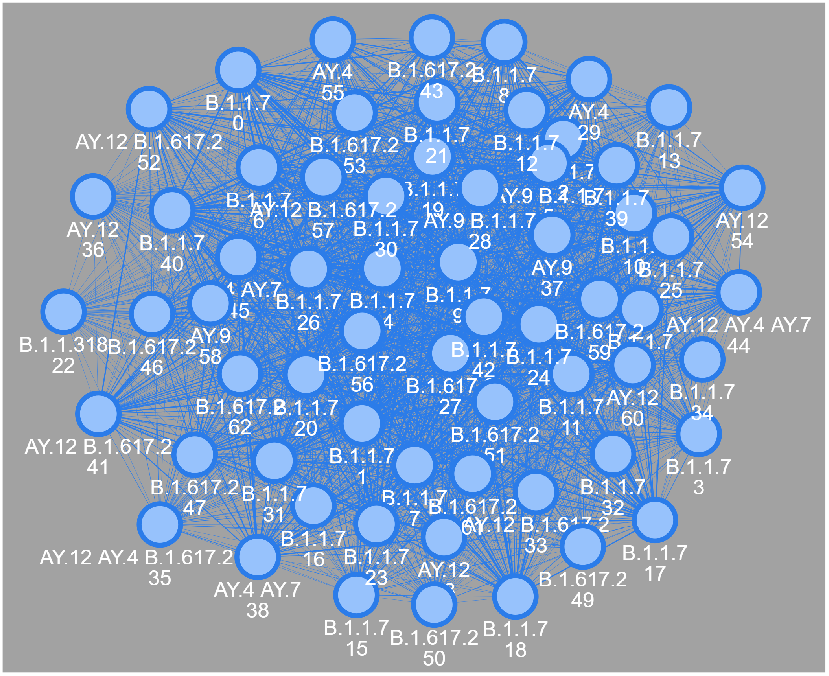
Undirected graph. MCL clustering has not yet been applied to that graph. One can easily see that it is almost fully connected. Underneath each node is its unique id and the assigned lineages to its samples. The nodes are connected to each other according to algorithm 1.

**Fig. 10.**
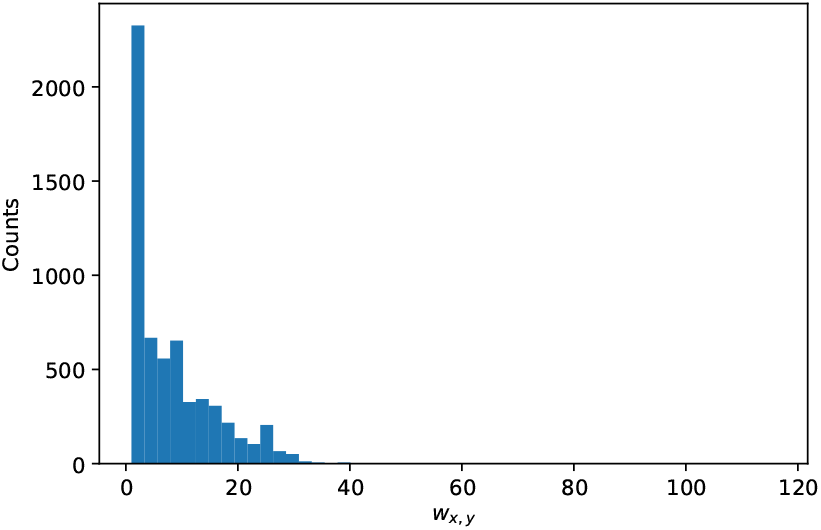
Histogram of graph’s weights *w_x, y_* prior to the mathematical transformation. The entire range of values of the distinct variable *w_x, y_* is divided into a series of nonoverlapping intervals (bins) of arbitrary length for better visualization. One could see that the vast majority of the weights is gathered between 0 and 32-ish. Note that the reason why it seems like there are no weights (counts = 0) with values greater than 32 is due to the fact that the number of weights with values greater that 32 is negligible.

Let

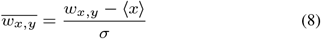

be the standardize weights, where:

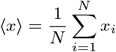

is the arithmetic mean, and:

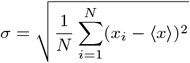

is the population standard deviation of a discrete random variable by using summation notation.

Given the Normal Distribution Probability Density Function (PDF)

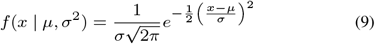

with mean *μ* and variance *σ*^2^ the, Normal Distribution Cumulative Density Function (CDF)

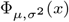

is calculated by the integral of equation 9. Those integrals cannot be expressed in terms of elementary functions, and are often claimed to be special functions. However, many numerical approximations are known which are beyond the scope of this thesis and that is why they are not shown.

Let

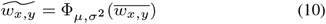

be the normalized weights in terms of transforming the standardized weights in such a way that they follow the Normal CDF. The distribution of those mathematically transformed weights 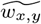 is shown in Fig. 11. We ended up using 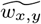 accompanying each pair of nodes {*x, y*} as the new input in the MCL algorithm.

**Fig. 11.**
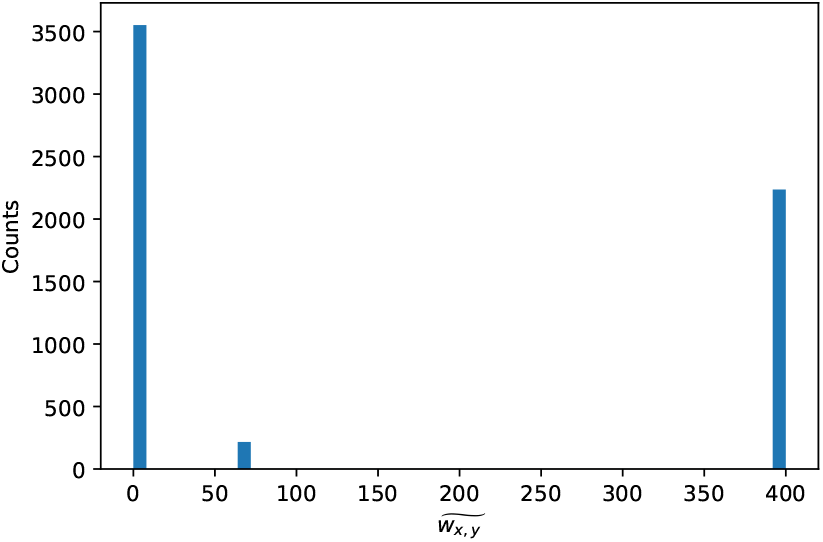
Histogram of graph’s weights 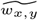 post the mathematical transformation. The entire range of values of the distinct variable 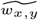 is divided into a series of non-overlapping intervals (bins) of arbitrary length for better visualization. One could point out a few key points. Firstly, all weights are now more spread out than they were before the transformation. Those that used to be closer to 0 have collapsed to 0 while those that used to be closer to the maximum value have collapsed to the new maximum value. Secondly, all weights are now bounded from a minimum allowed value of 0 to 400, which according to MCL documentation is an appropriate range.

The decision of inflating the gaps and stretching the distribution of weights in figure 10 through the method discussed in this chapter results in MCL’s distinctive ability to form highly interconnected clusters of nodes having increased dramatically. Since that method is an unsupervised clustering technique, there is no way to validate the results. On the other hand, one could use the percentage content of the compound lineages of each node of the graph in order to somehow validate the MCL’s clusters. To keep things as simple as possible we took into consideration only those nodes that consist of a dominant lineage with a percentage content of at least 90% or more. What we concluded is that nodes consisting of the same dominant lineages are gathered into distinct MCL clusters, which strongly indicates that the MCL algorithm is performing well on our dataset after the discussed transformation of the weights.

### 4.7 Arrow edges and co-occurring mutations

Co-occurring mutations in terms of positions of concern in common have already been determined through the methodology of chapter 4.5. Although these positions of concern have been providing insightful information for grouping different nodes of samples, they have not been providing any useful information regarding the evolution of the virus at all. On the other hand, an algorithmic procedure that identifies, between two different nodes, an actual mutation appearing in only one node and not in the other, in the same position of concern could potentially be used in order to draw an arrow from the node the mutation is missing to the node the mutation is popping up. As explained later on this chapter in detail, these arrows could be used to construct a directed graph indicating evolutionary paths of the virus.

#### Nodes

Given *A*_1_ and *A*_2_ the collection of the unique clusters’ identifiers based on 1s (non-characteristic mutations) and on 2s (characteristic mutations), we define, for each *a*_1_ ∈ *A*_1_ and for each *a*_2_ ∈ *A*_2_, the following sets:

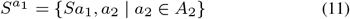

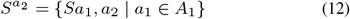

*S*^*a*_1_^, *S*^*a*_2_^ are sets of nodes given a fixed *a*_1_ or *a*_2_ respectively. We need to remind here that each node is the set of samples *S*_*a*_1_, *a*_2__, as defined by equation 3 for a given pair (*a*_1_, *a*_2_) ∈ *A*_1_ × *A*_2_.

##### Example 4.4.

*Given a collection of nodes* {*S*_0,2_, *S*_1,2_, *S*_3,0_} *each of which is a subset of S as equation 3 defines and assuming that each of those nodes is not an empty set, equation 11 will give:*

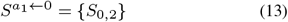

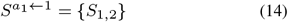

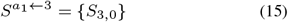

*and the equation 12 will give:*

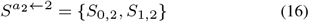

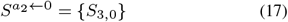

In other words, keeping the number of cluster fixed either based on “1s” or on “2s”, equation 11 will give us the group of nodes that belongs to a fixed cluster *a*_1_ based on “1s”, regardless of which clusters those nodes belong to based on “2s” and vice versa. Equation 12 will give us the group of nodes that belongs to a fixed cluster *a*_2_ based on “2s”, regardless of which clusters those nodes belong to based on “1s”.

As we move forward in this chapter, we will be referring to those sets as *S*^*a*_1_^ and *S*^*a*_2_^ of equations 11, 12 implying that we have already fixed the cluster *a*_1_ or *a*_2_ respectively. The example 4.4 demonstrates exactly this behavior.

#### Directed edges

Algorithm 2 is used to calculate arrows between pairs of nodes. An arrow is drawn from node *x* to node *y* if they have at least one PoC in common and that PoC is unmutated along the samples of the former node and mutated along the samples of the latter node. In that case node *x* becomes the starting point of the arrow while node *y* becomes its ending point.

Not all possible pairs of nodes are candidates for that method. More specifically, only those nodes that belong to a group 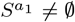 for a fixed *a*_1_ ∈ *A*_1_ are allowed to be connected to each other. Respectively, only those nodes that belong to a group 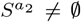 for a fixed *a*_2_ ∈ *A*_2_ are allowed to be connected to each other. If an arrow is drawn between a pair of nodes which belongs to a group given by equation 11, then this arrow is marked as of blue color and corresponds to the popping up of a non-characteristic mutation moving from one node to another. Alternately, if an arrow is drawn between a pair of nodes which belongs to a group given by equation 12, then this arrow is marked as of red color and corresponds to the popping up of a characteristic mutation moving from one node to another.

Having a closer look at example 4.4, it is made clear that only the nodes belonging to a specific group which is given by one of the equations 13–17 are allowed to be connected to each other. For instance, nodes *S*_0, 2_, *S*_1, 2_ of equation 16 are allowed to be connected to each other with a red arrow yet they are not allowed to be connected to any other nodes of equations 13, 14, 15, 17. In this particular example only nodes of equation 16 can be connected to each other simply because they are two. All the rest of the groups cannot form arrows among their nodes due to the fact that they consist of only one node. It becomes quite obvious that an arrow can be drawn if and only if there are enough nodes to be connected to each other inside a group. In other words the following constraint must be imposed:

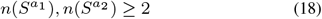

We define:

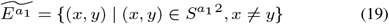

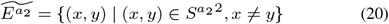

two sets of all plausible edges (also called directed edges, directed links, directed lines, arrows or arcs) which are ordered pairs of nodes (that is, an edge is associated with two distinct nodes). 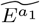 consists of all possible blue edges taking into consideration only non-characteristic mutations (“1”s), while 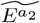 consists of all possible red edges taking into consideration only characteristic mutations (“2”s). Each ordered pair (*x, y*) represents a directed edge or an arrow, from node *x* to node *y*. Nodes *x* and *y* are called the endpoints of the edge, *x* the tail of the edge and *y* the head of the edge. The edge (*y, x*) is called the inverted edge of (x, y) and it is permitted. Multiple edges, not allowed under the definition of equations 19, 20, are two or more edges with both the same tail and the same head. Loops, edges that join nodes to themselves are not permitted since a directed loop would be (*x, x*) which is not in either 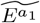 or 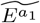 under the constraint *x* ≠ *y*.

Lines 1-22 and 24-45 of algorithm 2 are responsible for selecting two subsets of blue and red directed edges 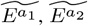 based on “1s” and “2s” respectively. In reality, these two blocks of code were implemented inside a for loop running twice, one regarding the based on “1s” implementation and one regarding the based on “2s” implementation. The selected blue directed edges are stored in the 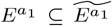 subset while the selected red directed edges in the 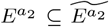 subset, as records in the form of ordered pairs (*x, y*).

The key points of the algorithm 2 are those on lines 11, 13 for blue edges and 34, 36 for red edges respectively. By iterating through all combinations of nodes (*x, y*) in *S*^*a*_1_^ or *S*^*a*_2_^ and consequently, through each PoC of node *x* and *y*, we check whether:

- the number of SNPs in those PoCs of *x* is zero (lines: 11, 34);
- the number of SNPs in the PoCs of *y*, which are in common with those of node *x*, is greater than zero (lines: 13, 36).

If the above-mentioned criteria are fulfilled, a new ordered pair (*x, y*) is added in *E*^*a*_1_^ or *E*^*a*_2_^ (blue or red directed edge), respectively.

Miscellaneous calculations are not shown in algorithm 2. In reality, additional data accompanying each directed edge in *E*^*a*_1_^ or *E*^*a*_2_^ are calculated as well. For instance, the specific SNPs and the number of each SNP’s copies in each PoC of the pointing node *y*, are composed in a table unique for each directed edge (*x, y*), as it is shown in example 4.5.

#### Directed graph definition

Having defined *S*^*a*_1_^, *S*^*a*_2_^ (see eq. 11, 12) and constructed *E*^*a*_1_^, *E*^*a*_2_^ through algorithm 2, as subsets of 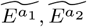 (see eq. 19, 20), let:

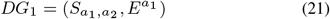

be a directed graph, making use of blue edges and:

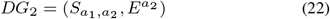

be a directed graph, making use of red edges.

In other words, tuples comprising:

- *S*_*a*_1_, *a*_2__, a set whose elements are called vertices (or nodes);
- *E*^*a*_1_^ or *E*^*a*_2_^, sets of ordered pairs of nodes, called directed edges.

Despite the fact that the *S*_*a*_1_, *a*_2__ and the *E*^*a*_1_^, *E*^*a*_2_^ are used in order to define these two graphs (see eq. 21, 22) and, by definition, these sets consist of groups of samples in the case of *S*_*a*_1_, *a*_2__ and directed edges connecting two elements of *S*_*a*_1_, *a*_2__ at a time in the case of *E*^*a*_1_^, *E*^*a*_2_^, in reality, as it has already been mentioned, both the nodes and the edges are accompanied by additional supplementary information.

A small table accompanying each node is calculated and includes exactly the same supplementary information as of those described in section “Graph definition” of chapter 4.5.

As for the directed edges, they are accompanied by a corresponding table such as those that have already been mentioned in example 4.5. For visualization purposes, the directed edges are visualized with colored arrows, blue in the case of *DG*_1_ graph and red in the case of *DG*_2_.

There are two different directed graphs, *DG*_1_ and *DG*_2_. The reason why two different directed graphs were defined is the fact that, in the case of one directed graph making use of both blue and red edges, there could have been multiple edges from one node to the other. Unfortunately, multiple edges are not permitted by the definition of the directed graph. In a more general sense, a graph allowing multiple edges, could be defined, yet things have been kept as simple as possible. Instead, two different simple directed graphs, one making use of blue directed edges and one making use of red directed edges were used. Since the exact same nodes were used in order to define both graphs, we were able to visualize them together, the one on top of each other in a way that seemed to be a single directed multigraph, figure 12.

**Fig. 12.**
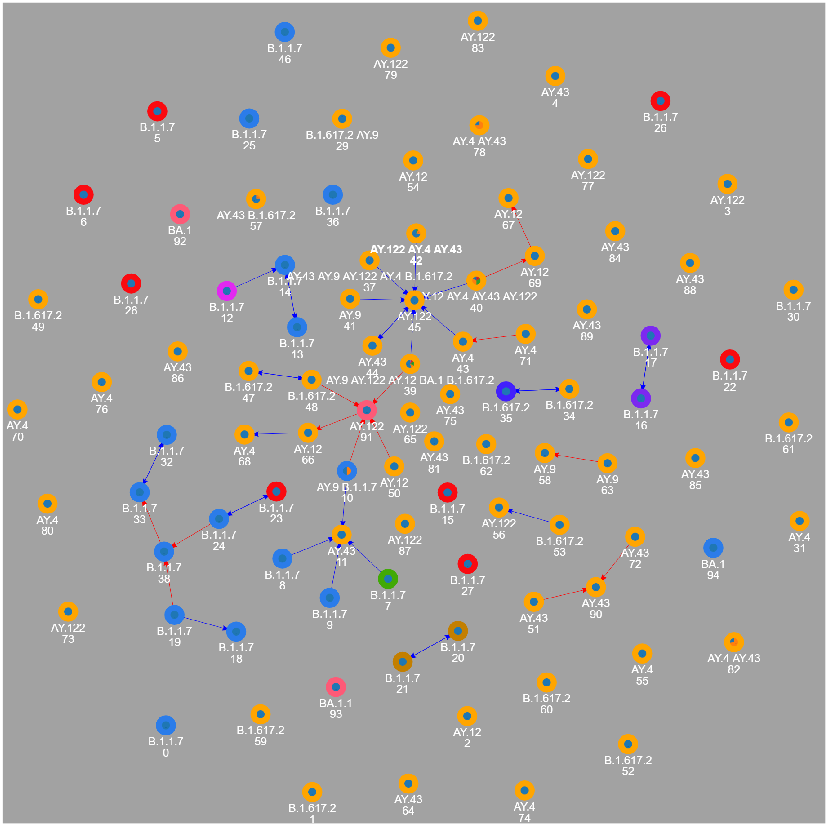
Directed graph of two networks, with the one plotted on top of each other as it was a single directed multigraph with two types of edges, blue for mutations found analyzing clusters formed by non-characteristic mutations and red for mutations found analyzing clusters formed by characteristic mutations. Each node’s outer color (colorful circular ring) corresponds to the high interconnectivity group that each node has been assigned to by the MCL. The pie chart inside the circular ring is the percentage content of the compound lineages that node is made of while the compound lineages by themselves are mentioned in the first line underneath each node. The second line underneath the nodes is a unique identifier of the corresponding node.

### 4.8 Multiple sequence alignment (MSA)

In section 4.7, we defined a graph, the edges of which connect nodes to each other in a useful way, as a tool of finding groups of samples closely related to each other, and identifying mutations popping up from one group of samples to another, in an attempt to gain an understanding of the evolution of SARS-CoV-2. As a matter of course, it is if not impossible, extremely difficult for one to determine the evolutionary history of the virus or “who infected whom” [22]. For that reason and just because, as far as we are aware, at the time of writing, there is no existing methodology capable of addressing that problem, we will try to validate the results obtained by the graph analysis and its directed edges by using a multiple sequence alignment as explained below.

By obtaining an accurate multiple sequence alignment of approximately 4.500 SARS-CoV-2 genomes of 30.000-ish bases soon made it clear that it is a tremendously computational expensive task. It turns out that by employing the rapid calculation of full-length MSA of closely-related viral genomes mode of MAFFT aligner (cite MAFFT), a full-length MSA of our SARS-CoV-2 genomes is a relatively computational easy task in terms of a few minutes on a regular PC. It is worth mentioning here that according to MAFFT documentation it is sometimes useful and vastly faster to align all sequences just to the reference genome to build a full MSA. However, by doing so, it is very likely for gaps to be inserted into the reference sequence if deemed necessary hence altering its initial length. To avoid this behavior *keeplength* setting was used since it reassures no gaps are inserted into the reference sequence. Furthermore, *maxambiguous* was set to 1 to prevent removing sequences that have ambiguous letters while *ep* or offset value was set to 0.1 as it’s been recommended if no long gaps are expected.

#### Example 4.5.

*Given an ordered pair* 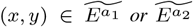, *which represents a posible directed edge connecting nodes x and y, the following table is their content:*

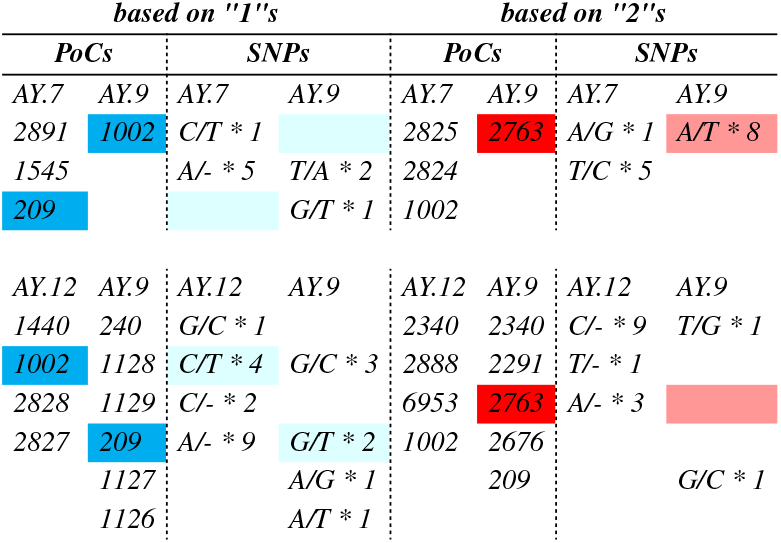

*Where the first group of samples corresponds to node x, while the second one corresponds to node y. It is worth mentioning that there might be the case where more than one SNP have been found in a PoC. If that happened, for example, the new record would be “G/C * 1, A/T * 3”. The PoCs in common between nodes x, y based on “1”s and “2”s are:*

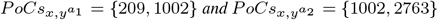

*The number of SNPs found in the common PoCs* 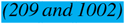 *between nodes x and y, is* 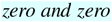 *for x and* 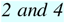 *for y, respectively. Since there is at least one SNP popping up from node x* → *y, in the same PoCs, under the -based on “1”s-analysis, a directed edge will be drawn from x to y and the following table accompanying that edge will be:*

| SNPs | counts | position |
| --- | --- | --- |
| G/T | 2 | 209 |
| C/T | 4 | 1002 |

*and*

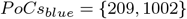

*The number of SNPs found in the common PoCs* 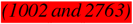 *between nodes x and y, is* 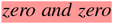 *for y and* 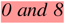 *for y, respectively. Since there is at least one SNP popping up from node y* → *x, in the same PoCs, under the -based on “2”s-analysis, a directed edge will be drawn from y to x and the following table accompanying that edge will be:*

| SNPs | counts | position |
| --- | --- | --- |
| A/T | 8 | 2763 |

*and*

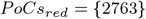

***Each data mentioned in this example is just dummy values and does not represent any real information.***

#### Algorithm 2 Calculate directed edges

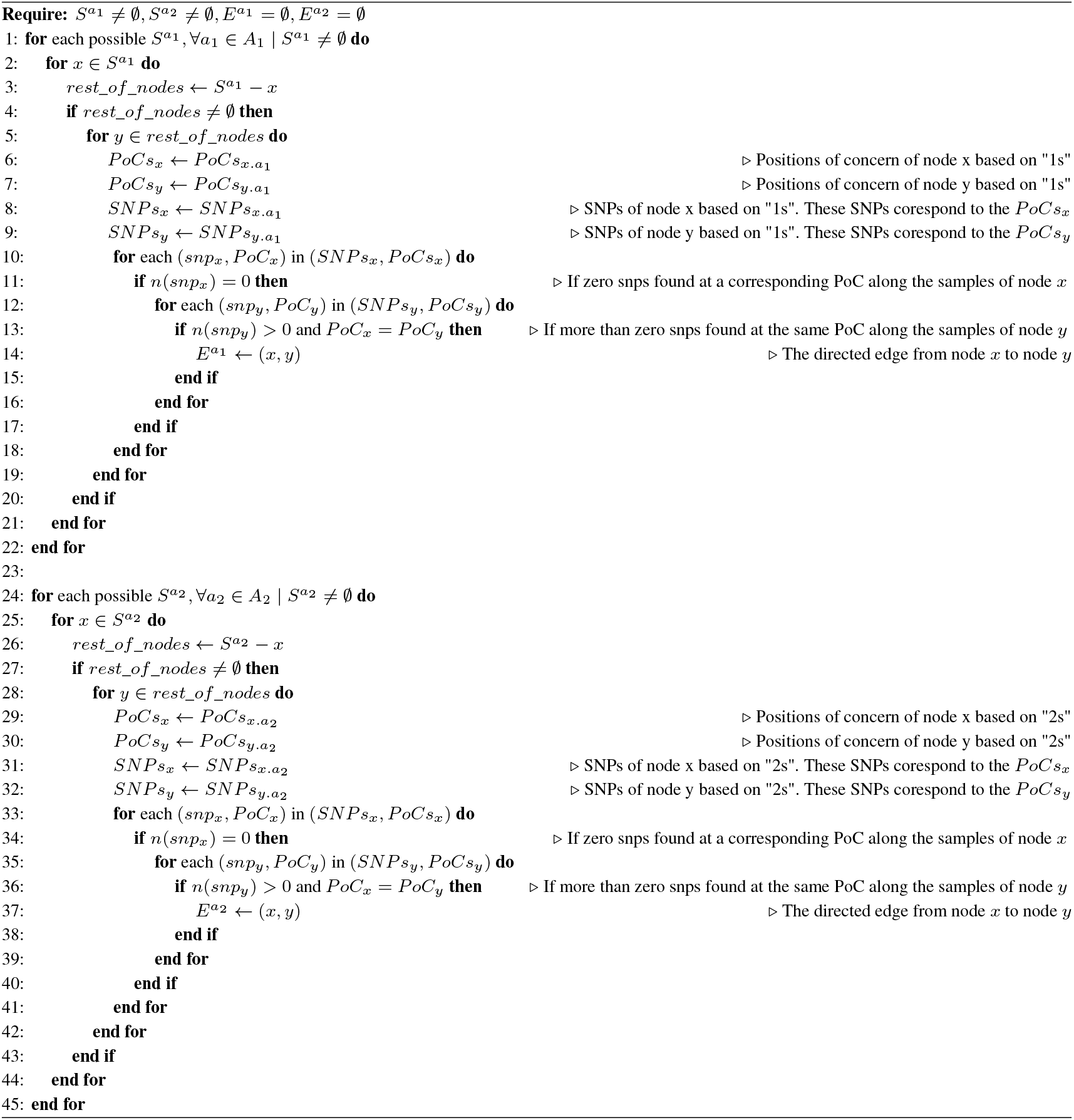

The rapid calculation of full-length MSA of closely-related viral genomes mode of MAFFT aligner was originally for other purposes, yet the SARS-CoV-2 outbreak made clear that this method can be used efficiently and effectively on SARS-CoV-2 genomes, and it has become widely known in the community ever since. Placing trust in the community’s wide usage of it on SARS-CoV-2 data we have gone through this method without validating the MSA by any means.

### 4.9 Entropies - Mutual information gain

Having a closer look at graph 12, one can identify strongly connected groups of nodes with either blue or red or even both types of edges. One can even move/rearrange the nodes through the drag and drop mode of the used GUI, making it even easier for the human eye to point out a few groups of interest, resulting in the graph 15. Once a particular group of nodes has been identified (see graph 15 selected areas), the nodes ids of that specific group are needed as input for the next step. The samples that each node of interest is made of are then obtained. Following the same notation, we will be calling those nodes, nodes of interest and their samples, samples of interest. Given the full MSA comprised of all samples and the samples of interest, we are slicing the former obtaining only a part of it that consists only of the samples of interest, while maintaining all its required properties. At the same time, we obtain the reported positions by all directed edges, blue and red, that connect the nodes of interest to each other, and we store them in a suitable data structure. From now on we will be referring to those positions as sites of interest.

Possessing an MSA of only the samples of interest and their sites of interest provided by the edges that connect nodes of interest, we calculated the mutual information (MI) of each possible pair of the abovementioned sites of interest.

Let

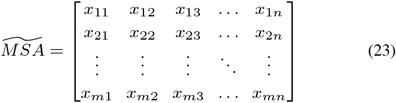

be the slice of the full MSA with each *x_i,j_* representing the *j*^th^ base/site/position of the *i*^th^ sample.

Let also *PoCs_blue_*, *PoCs_red_* be the sites of interest that have been reported by the total number of blue and red edges, respectively, connecting nodes of interest (In example 4.5 there are only two edges, one blue and one red. *PoCs_blue_* and *PoCs_red_* would represent the position columns of the two accompanying tables. In the case of a group of nodes of interest, considering more blue and red edges, *PoCs_blue_* would be the union of all blue position columns of the accompanying tables reported by blue edges while *PoCs_red_* would be the union of all red position columns of the accompanying tables reported by red edges).

In order to obtain the total reported positions by the total blue and red edges in the group of nodes of interest, we take the union between the reported positions by blue edges and the reported positions by red edges. We then have

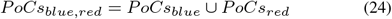

Equation 24 gives the total positions of interest or, in other words, some columns of the 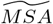 that we are interested in. Then the

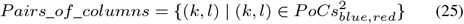

consists of all posible combinations in pairs of two columns, given by the cartesian product of *PoCs_blue, red_* with itself.

Considering the columns *PoCs_blue, red_* as random variables, the probabilities of each character in a random variable (column), *x_j_* ∀*j* ∈ *PoCs_blue, red_*, along the *m* samples of the 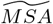 are given by the count of each character over the total counts of all different characters given by the random variable. Since, in our case, the length columns is constant and equals to *m*, and the value of a random variable can be one of *A, T, G, C*, the list of probabilities for a particular random variable/column is given by

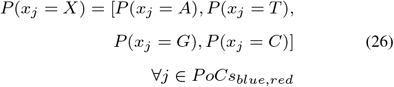

where

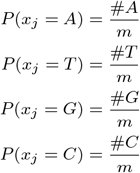

Here the symbol # is used to denote the number of occurrences. Due to the fact that 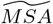 is a multiple sequence alignment, there could be spaces among the sites of different samples and thus, the “-” symbol should be included in the counts. For simplicity reasons though, we have not included it in our analysis.

Having obtained a part of the full MSA that is related to the group of nodes of interest, the next logical step is to use a metric in an attempt to measure the dependency of the sites of interest in that sub-MSA. For that purpose the mutual information is acquired. Mutual information (MI) is non-negative value, which shows the dependency between two variables. In our case the random variables are discrete random variables. The higher the MI value is, the more dependent the variables are. The normalized MI is then calculated by using the `“normalized_mutual_info_score”` method of scikit learn library, between all pairs of the discrete random variables/columns, *Pairs_of_columns*, given in pairs of two by equation 25. NMI is a normalization of the MI score. The result is a square *d* × *d* matrix, where *d* = *n*(*PoCs_blue, red_*) is the cardinality of *PoCs_blue, red_*, and its values are scaled from 0 to 1. 1 shows perfect correlation while 0 means no correlation at all. That is because the NMI was calculated bewteen all pairs of the sites of interest given by *PoCs_blue, red_* and hence, the number of elements of that set is the number of unique variables. Figure 13 shows a heatmap of the calculated NMI between all pairs of sites of interest in a group of nodes of interest.

**Fig. 13.**
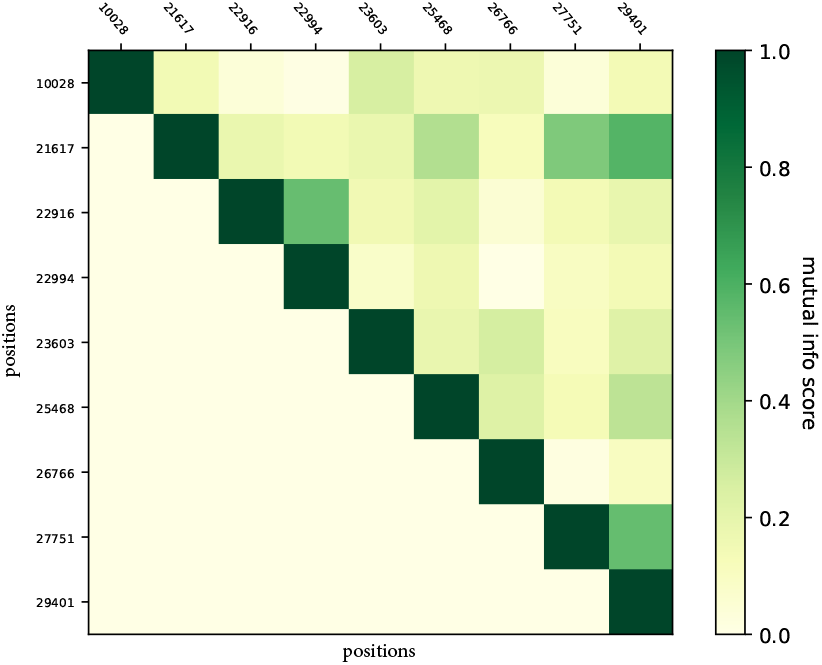
Heatmap of the Normalized Mutual Information (NMI) between all pairs of sites of interest in the group of nodes of interest 39, 50, 10, 48, 66, 91 indicated by the directed graph 12. X and y axis show the sites of interest. The darker the color the more closely related the corresponding sites are. All on-main-diagonal elements are one due to the fact that it is the NMI calculated between a site and its itself. Only the upper triangular part was needed to be calculated because this heatmap is square and completely symmetrical, saving a little bit of computational time.

Hierarchical clustering on the sites of interest is then performed (figures: 17, 16, 14, 19, 18, 20). Specifically, the “AgglomerativeClustering” method of scikit learn library, with the default parameters is used, except for linkage wich is set to “complete”. Agglomerative clustering needs a distance metric. By definition, a lower distance is better than a higher one and vice versa. In the case of the NMI values, though this criterion is not fulfilled. On the contrary, NMI behaves in exactly the oposite way, a higher NMI value corresponds to a higher dependency while a lower NMI to a weaker dependency. For that reason, we reverse the NMI values, using the following formula

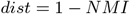

transforming the NMI values to a distance metric, where, a lower value, close to 0 means that the in comparison variables are highly correlated and hence are closely related to each other, while, a high distance, close to 1, means that the variables are weakly correlated and, thus, they are very distant to each other. It is worth mentioning that due to rounding errors negative distances may occur. In that case, they are set to zero, since 0 is the lowest possible value.

**Fig. 14.**
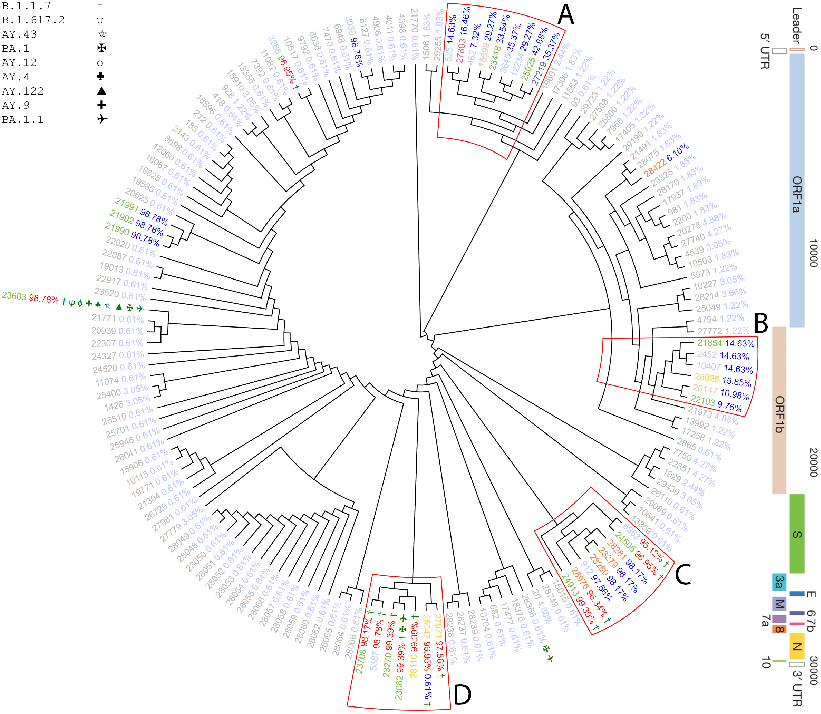
Hierarchical clustering of the reversed Normalized Mutual Information between all pairs of sites reported by the directed graph 12, taking into account nodes 32, 33, 38, 24, 23, 19, 18 (number of samples used = 164). Reading each leaf of the dendrogram from left to right, one can descry at most 4 fields of information. 1) A black or colorful number which represents a position/location/site of interest, the color of which shows the gene that belongs to (see vertical bar). 2) A percentage colored in red is shown, indicating the proportion of samples in which this mutation has been identified as characteristic over the total number of the occurrences of that mutation along the samples of concern. 3) A percentage colored in blue is shown, indicating the proportion of samples in which this mutation has been identified as non-characteristic over the total number of the occurrences of that mutation along those samples of concern. Notice that, the red percentage is not shown when this mutation has not been identified as a characteristic mutation, while the blue percentage is not shown when this mutation has not been identified as a non-characteristic mutation. For instance, a mutated position that has been found as characteristic mutation in half of the samples of concern and non-characteristic in the other half of the samples, would have 50-50% red and blue percentages. 4) The following green symbols indicate the lineages of the samples in which this mutation has been found (see the upper-left corner for the available lineages). The black or faded positions/locations/sites are considered insignificant and correspond to mutations that have been identified as non-characteristic mutations at less that 5% and (not) as characteristic mutations at 0%.

**Fig. 15.**
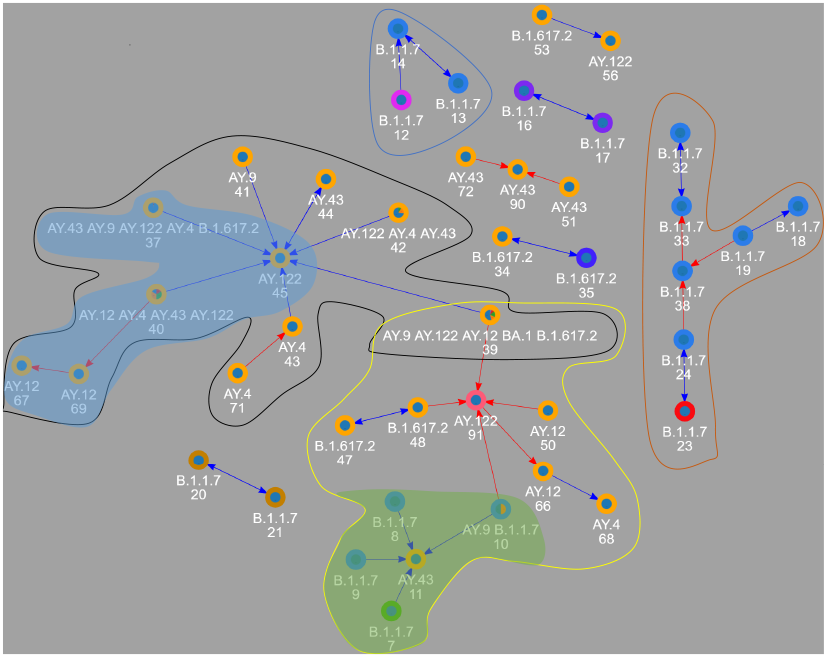
Part of the graph 12 that could potentially indicate interesting evolutionary paths of the virus. Human choice is shown using either colored closed loops or shadowed areas for simplicity reasons due to overlapping areas.

**Fig. 16.**
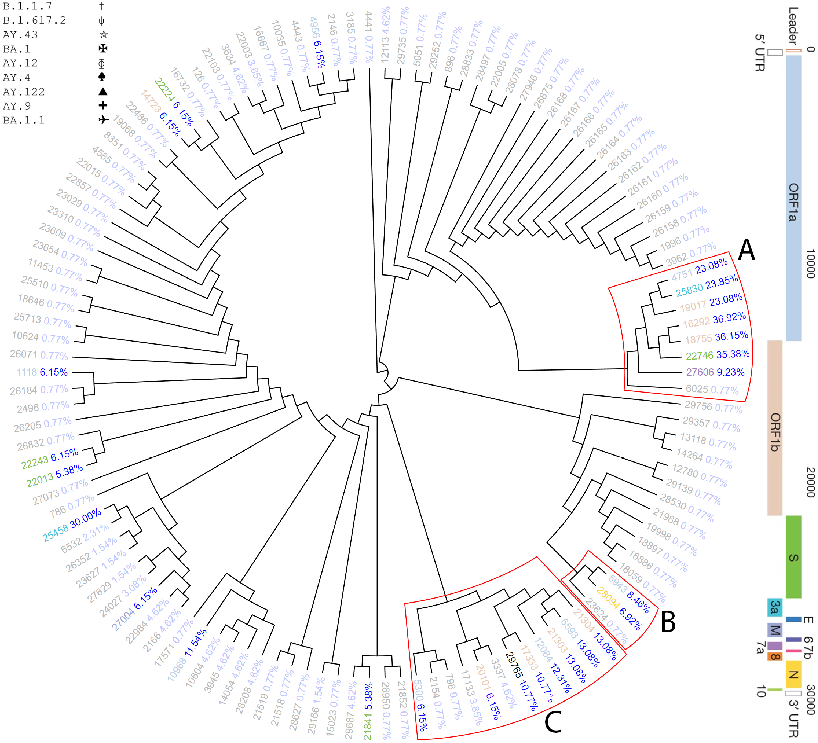
Hierarchical clustering of the reversed Normalized Mutual Information between all pairs of sites reported by the directed graph 12, taking into account nodes 12, 13, 14 (number of samples used = 130).

**Fig. 17.**
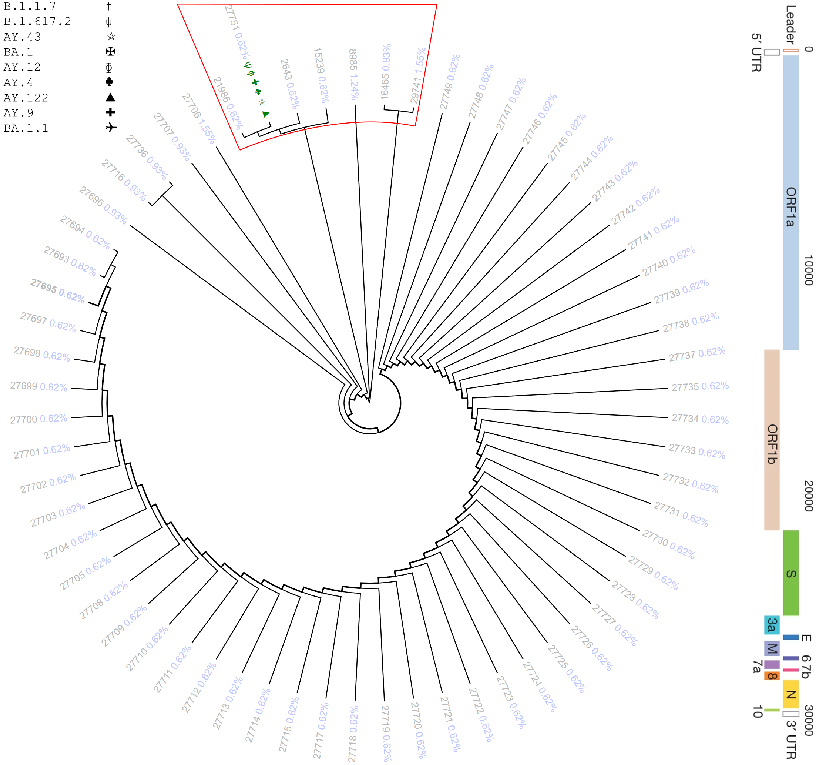
Hierarchical clustering of the reversed Normalized Mutual Information between all pairs of sites reported by the directed graph 12, taking into account nodes 10, 7, 9, 8, 11 (number of samples used = 323).

**Fig. 18.**
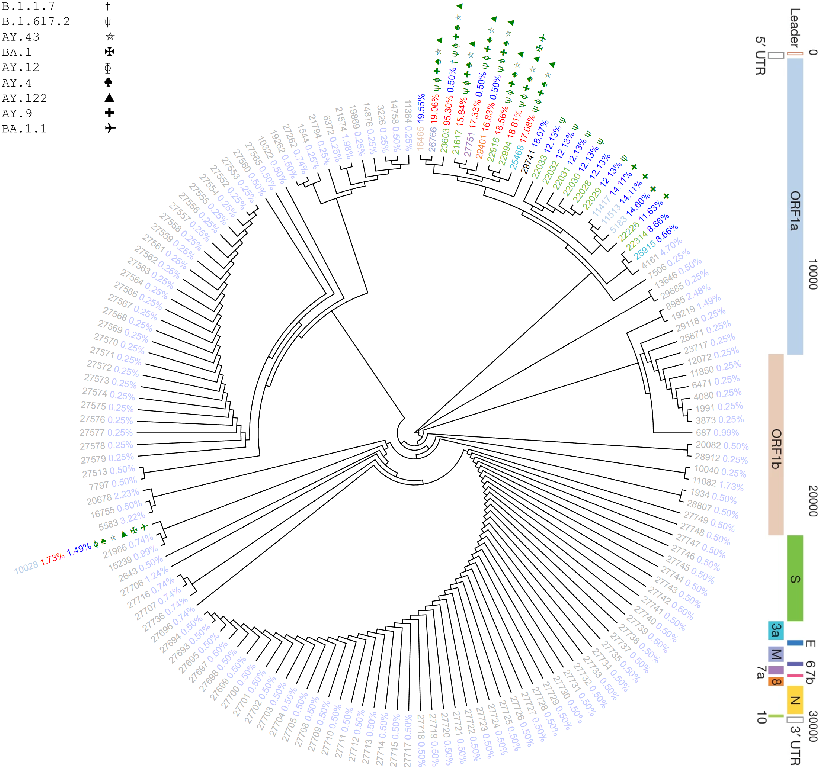
Hierarchical clustering of the reversed Normalized Mutual Information between all pairs of sites shown in directed graph 12, taking into account nodes 39, 47, 48, 91, 50, 66, 68, 10, 11, 7, 9, 8 (number of samples used = 404).

**Fig. 19.**
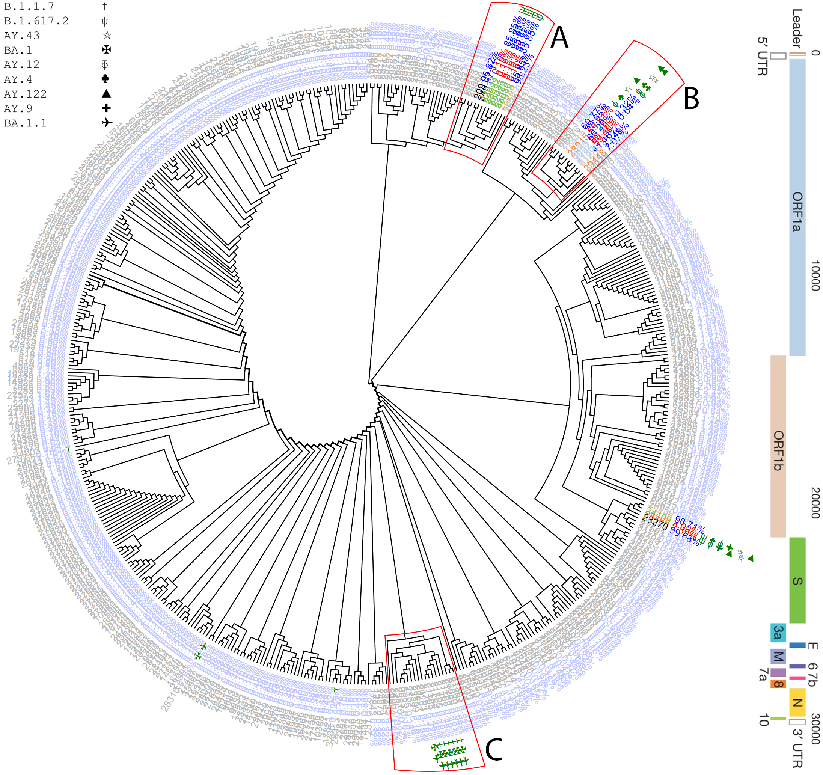
Hierarchical clustering of the reversed Normalized Mutual Information between all pairs of sites reported by the directed graph 12, taking into account nodes 37, 45, 40, 69, 67 (number of samples used = 622).

**Fig. 20.**
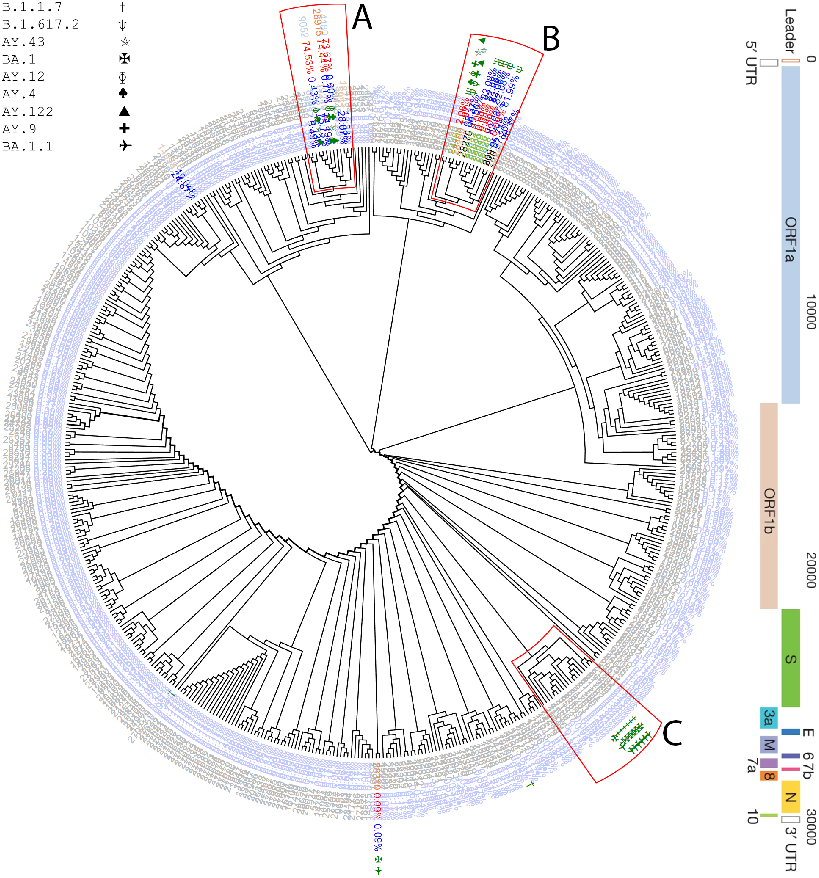
Hierarchical clustering of the reversed Normalized Mutual Information between all pairs of sites reported by the directed graph 12, taking into account nodes 39, 71, 43, 45, 42, 44, 41, 37, 40, 69, 67 (number of samples used = 1166).

## 5 Results

Conducted studies have shown that many mutations can randomly occur in the time of virus replication inside host’s body [25, 21]. An enormous number of factors, both environmental and from inside host’s body, could potentially cause the necessary selective pressure, which can result in some random mutations being selected. Going through our methodology, we obtained graph 15 which could potentially indicate evolutionary paths of the virus, among well known lineages, revealing randomly co-occuring mutations beyond strain-specific / strain-defining ones. Graph 15 is a small part of the whole graph 12 which is the result of algorithm 2. After careful consideration, we identified 6 distinct groups of nodes of interest shown as either colored areas or inside colored curves in graph 15. For each of these groups of nodes, a dendrogram is provided, as a result of the hierarchical clustering of the reversed Normalized Mutual Information (see chapters 4.8 and 4.9).

For instance, nodes 32, 33, 38, 24, 23, 19, and 18 from inside the orange curve constitute a homogeneous network consisting of samples that have been classified as of lineage B.1.1.7. Important and strongly correlated sites among these samples are shown in figure 14. The small sub-network inside the blue closed loop that is comprised of nodes 12, 13, and 14 is a small homogeneous network consisting of samples that have been classified as of lineage B.1.1.7. Its sites of interest are shown in figure 16. The big sub-network inside the yellow and black closed loops has been split into two different parts, maintaining an overlap of one node, since the sites/positions of concern were so many that it would be very difficult for one to interpret the final dendrogram. The sub-network inside the yellow area that consists of nodes 39, 91, 48, 47, 50, 66, 68, 10, 11, 7, 8, and 9, is highly inhomogeneous containing samples of the European lineage, Alpha, Delta, and Omicron variants. The hierarchical clustering of important and strongly correlated sites found among its samples are shown in figure 18. By disassembling the sub-network inside the yellow area even more into its constituent pieces, we conclude that the sub-network in the green shadowed area is also of high interest. It is assembled of 7, 8, 9, 10, 11 nodes and it consists mainly of B.1.1.7 (Alpha variant) samples and a small number of AY.43, alias of the European lineage and AY.9, alias of the UK lineage (Alpha variant) (Significantly important sites are reported in figure 17). The sub-network inside the black loop that consists of nodes 39, 45, 42, 44, 41, 37, 40, 69, 67, 43, and 71, is very inhomogeneous and indistinguishable in terms of lineages which makes it worth the investigation. There are many lineages, such as, AY.9 (Alpha variant), AY.122 and AY.43 (European lineage), BA.1 (Omicron variant), and B.1.617.2 (Delta variant) (see figure 20). Furthermore, taking apart the sub-network inside the black closed loop into its constituent pieces, we figured out that the sub-network within the blue shadowed area needs to be further investigated. It is comprised of nodes 45, 37, 40, 69, and 67 with high rates of EU and UK lineages and Delta variants (figure 19).

## 6 Discussion & Conclusions

In this study, we first attempted to cluster the available SARS-CoV-2 samples in groups, and to validate those groups based on the reported characteristic mutations by the pangolin tool [15]. Then co-occurring mutations among these groups were counted and further revealed the phylogenetic relationships among groups of samples with either coexisting or by counting the occurrences of important mutations moving from one group to the other. Our methodology was validated by calculating the Mutual Information between pairs of sites in a multiple sequence alignment, along samples indicated by the previously mentioned groups. We then used the mutual information in a way to represent the distance of those sites from each other. We ended up performing hierarchical clustering on those distances obtaining potentially useful insights regarding the evolutionary history of the virus and validating the phylogeny among our groups.

Specifically, figure 14 shows that identifying evolutionary patterns among B.1.1.7 samples with known labels could be possible. Based on the non-characteristic mutated sites at medium to high rates of group A and the fact that this group is close to the root of the tree, we conclude that evolutionary pressure could lead to new B.1.1.7 sub-lineages, forcing those mutations to prevail. We can draw the same conclusions based on group B yet by a weaker indication since group B’s sites appear to be mutated at lower percentages. On the other hand, the high prevalence of non-characteristic and characteristic mutations that belong to B.1.1.7 lineage, at very high rates, in group C could indicate the circulation of non-characteristic mutations closely related to characteristic mutations revealing useful patterns. Moreover, the presence of a B.1.1.7-characteristic mutation (23062) at very high rates that belongs to not only the B.1.1.7, but also to the BA.1 and BA.1.1 and the fact that it is strongly correlated to other B.1.1.7-characteristic mutations, could help us identify potentially important mutations in feature lineages, such as, BA.1 and BA.1.1 of omicron variant.

In figure 17 the majority of the reported sites appears to be insignificant non-characteristic mutated at very low rates and thus figure 17 is of less importance. However, the first 3 branches, starting from the root of the tree, are the most remote groups of mutations and hence they are worth investigating. The mutation at position 27751, is a non-characteristic mutation along the samples used to construct this tree, appearing at very low rates. Yet, it is a characteristic one when it is studied in lineages B.1.617.2, AY.12, AY.9, AY.4, AY.43, AY.122. It is worth mentioning that samples of the above-mentioned lineages appear at higher frequencies as we move outside the green shadowed area to the rest of the nodes in the yellow area of figure 15, from node 10 to 91 and beyond. As a result, based on the existing labels we draw the conclusion that this mutation is very likely to pop up in future lineages.

In figure 18, one can clearly identify multiple non-characteristic and characteristic mutations that belong to alpha, beta, and delta variants at high rates, making this tree highly inhomogeneous. Surprisingly, there are some sites that while they are characteristic mutations of the compound lineages of this system, appear as non-characteristics at low rates. More specifically 22029-22033 and 22226, 5183, 11513, 11417 are reported as non-characteristic mutations while they are characteristic mutations of lineages B.1.617.2 and AY.9 respectively, lineages that are indeed included in the yellow area of figure 15. Additionally, some reported sites, such as 16465, 22314, and 25915 do not belong to any known lineage, at least at the time of writing, yet they were found as non-characteristic mutations along that system’s samples at significant rates. Those facts could again be indications of emerging mutations in future lineages or patterns between non-characteristic and characteristic mutations revealing the selective pressure of the former to the latter.

The tree in figure 16 is made of nodes 12, 13, 14 (graph 15) and is highly homogeneous since all of its used samples are B.1.1.7-assigned lineages. Interestingly, not even one site, reported by this tree appears as a characteristic mutation. This tree could potentially be used in order to identify emerging mutations of B.1.1.7 lineage or even to classify new samples to new sub-B.1.1.7-lineages. In groups, A B, and C one can observe non-characteristically mutated sites at middle to relatively high rates accompanying a few non-characteristically mutated sites at significantly low rates, such as 6025 in group A, 23624 in group B, and 2154, 798, 17133, 3337 in group C. That co-existence of non-characteristic mutations at middle to high and very low rates could be an indication of either new sub-variants that are being emerged or mutations that are faded out through the evolutionary history of the virus.

On the other hand, figures 19 and 20 are representative examples of how complex and indistinguishable the situation could become, if one included multiple nodes with a large number of samples in their analysis. There is a ton of characteristic and non-characteristic mutations in groups A and B of both trees since a tremendous number of samples have been included. It is worth mentioning that in group C of figure 19, non-characteristic mutations at insignificant rates pop up which, surprisingly, characterize lineages such as BA.1.1 and BA.1 that do not belong to the used part of the graph (blue shadowed area of graph 15). Unfortunately, not many conclusions can be drawn from these figures due to their complexity. Further analysis of these nodes must be done to safely draw conclusions regarding patterns between their characteristic and non-characteristic mutations.

Several studies attempting to either classify or even cluster SARS-CoV-2 populations have been suggested in the past. The majority of those methods depend on phylogenetic inference, bringing limitations regarding statistical errors, computation, and visualization in order for them to draw useful results. To avoid the excessively complex and computationally expensive phylogenetic construction, many novel sub-typing methods have been proposed, yet they are not able to determine the phylogenetic relationships among different sub-types [26]. Therefore, our proposed novel method replaces the classical phylogenetic approaches, reducing complexity while maintaining its ability to infer evolutionary paths of the virus. In addition to this, our method could update the phylogenetic model based on new-coming samples in a significantly more efficient and effective way, in a constantly evolving environment.

In our exploratory analysis, we assumed that the ground-truth labels, in order to validate the clustering analysis, are those assigned by the Pangolin system. Different labels of different nomenclature systems should be further used in future studies. It is also widely known that the dimensionally reduced space as a result of TSNE must not be used to perform clustering techniques on it, since the former distorts the distances among points to such an extent, that the results could be misleading. However, the fact that we validated the clusters based on the existing labels, lets us safely cluster the samples on the TSNE space despite those limitations. In other words, it was more of a supervised clustering than an unsupervised method.

Different modeling methods of the SARS-CoV-2 genomes could also be considered. For instance, instead of using the binary model we developed, one could use more sophisticated methods to encode those genomes into abstract mathematical representations. A particular use case of such embeddings is BioVec [1], which is a novel approach for representing biological sequences into vectors, through the application of unsupervised machine learning. This method has been evaluated in classifying proteins into protein families, achieving intriguing results. Therefore, one could use those mathematical representations (embedding vectors) in order to perform either direct clustering in that multidimensional space or dimensionality reduction followed by clustering. It is worth mentioning that, in this study, we attempted to use BioVec, yet due to computational limitations we drew the conclusion that it is beyond the scope of this manuscript. Further research must be done to examine whether such a model could be used for that purpose or could lead to better results. Different dimensionality reduction methods or clustering techniques could also be used, making sure that the clustering methods will be density based since that is a prerequisite of our methodology.

Here, we present a computational method for detecting patterns of co-occurring mutations potentially revealing the evolution of SARS-CoV-2. Not only did we identify emerging mutations that could lead to the emergence of new sub-linegages, but we also found non-characteristic mutations beyond strain-specific / strain-defining ones strongly correlated to strain characteristic mutations. Although these correlations might indicate an underlying “association” between different lineages, correlation does not necessarily mean causation. In order for one to identify causal relationships, more research is needed on drivers of evolution and the emergence of new mutations on perhaps worldwide data. As SARS-Cov-2 continues to spread globally, we hope our software will help detect new evolutionary events in the fight against the global pandemic.

## Code & Data Availability Statement

All SARS-CoV-2 sample data were retrieved from ENA (Project ID: PRJEB44141). All code developed for the analysis is available at https://github.com/BiodataAnalysisGroup/BEEMUS.

## Acknowledgments

We would like to acknowledge all the support and feedback we received from the Bioinformatics Laboratory at the Institute of Applied Biosciences at CERTH (INAB | CERTH).

## References

[1] E. Asgari and M. R. K. Mofrad. Continuous Distributed Representation of Biological Sequences for Deep Proteomics and Genomics. PLOS ONE, 10(11):e0141287, Nov. 2015.

[2] D. P. Bartel. Metazoan MicroRNAs. Cell, 173(1):20–51, Mar. 2018. Publisher: Elsevier.

[3] C.-E. Bichot. Population Based Metaheuristics, Fusion-Fission and Graph Partitioning Optimization. In P. S. Charles-Edmond Bichot, editor, Graph Partitioning, page 384. ISTE - Wiley, June 2011.

[4] P. Cingolani, A. Platts, L. L. Wang, M. Coon, T. Nguyen, L. Wang, S. J. Land, X. Lu, and D. M. Ruden. A program for annotating and predicting the effects of single nucleotide polymorphisms, SnpEff. Fly, 6(2):80–92, Apr. 2012. Publisher: Taylor & Francis _eprint: https://doi.org/10.4161/fly.19695.

[5] P. Danecek, A. Auton, G. Abecasis, C. A. Albers, E. Banks, M. A. DePristo, R. E. Handsaker, G. Lunter, G. T. Marth, S. T. Sherry, G. McVean, and R. Durbin. The variant call format and VCFtools. Bioinformatics, 27(15):2156–2158, Aug. 2011.

[6] S. M. v. Dongen. Graph clustering by flow simulation, May 2000. Accepted: 2001-02-13T10:26:00Z.

[7] M. Ester, H.-P. Kriegel, J. Sander, and X. Xu. A density-based algorithm for discovering clusters in large spatial databases with noise. In kdd, volume 96, pages 226–231, 1996. Issue: 34.

[8] G. E. Hinton and S. Roweis. Stochastic Neighbor Embedding. In Advances in Neural Information Processing Systems, volume 15. MIT Press, 2002.

[9] C. Huang, Y. Wang, X. Li, L. Ren, J. Zhao, Y. Hu, L. Zhang, G. Fan, J. Xu, X. Gu, Z. Cheng, T. Yu, J. Xia, Y. Wei, W. Wu, X. Xie, W. Yin, H. Li, M. Liu, Y. Xiao, H. Gao, L. Guo, J. Xie, G. Wang, R. Jiang, Z. Gao, Q. Jin, J. Wang, and B. Cao. Clinical features of patients infected with 2019 novel coronavirus in Wuhan, China. The Lancet, 395(10223):497–506, Feb. 2020.

[10] K. Katoh, K. Misawa, K.-i. Kuma, and T. Miyata. MAFFT: a novel method for rapid multiple sequence alignment based on fast Fourier transform. Nucleic Acids Research, 30(14):3059–3066, July 2002.

[11] R. P. Kincaid and C. S. Sullivan. Virus-Encoded microRNAs: An Overview and a Look to the Future. PLOS Pathogens, 8(12):e1003018, 2012. Publisher: Public Library of Science.

[12] L. Morales, J. C. Oliveros, R. Fernandez-Delgado, B. R. tenOever, L. Enjuanes, and I. Sola. SARS-CoV-Encoded Small RNAs Contribute to Infection-Associated Lung Pathology. Cell Host & Microbe, 21(3):344–355, Mar. 2017.

[13] B. Morel, P. Barbera, L. Czech, B. Bettisworth, L. Hübner, S. Lutteropp, D. Serdari, E.-G. Kostaki, I. Mamais, A. M. Kozlov, P. Pavlidis, D. Paraskevis, and A. Stamatakis. Phylogenetic Analysis of SARS-CoV-2 Data Is Difficult. Molecular Biology and Evolution, 38(5):1777–1791, May 2021.

[14] Á. O’Toole, V. Hill, O. G. Pybus, A. Watts, I. I. Bogoch, K. Khan, J. P. Messina, T. C.-. G. U. C.-U. Consortium, N. f. G. S. i. S. Africa (NGS-SA), B.-U. C. G. Network, H. Tegally, R. R. Lessells, J. Giandhari, S. Pillay, K. A. Tumedi, G. Nyepetsi, M. Kebabonye, M. Matsheka, M. Mine, S. Tokajian, H. Hassan, T. Salloum, G. Merhi, J. Koweyes, J. L. Geoghegan, J. d. Ligt, X. Ren, M. Storey, N. E. Freed, C. Pattabiraman, P. Prasad, A. S. Desai, R. Vasanthapuram, T. F. Schulz, L. Steinbrück, T. Stadler, S. V. S. Consortium, A. Parisi, A. Bianco, D. G. d. Viedma, S. Buenestado-Serrano, V. Borges, J. Isidro, S. Duarte, J. P. Gomes, N. S. Zuckerman, M. Mandelboim, O. Mor, T. Seemann, A. Arnott, J. Draper, M. Gall, W. Rawlinson, I. Deveson, S. Schlebusch, J. McMahon, L. Leong, C. K. Lim, M. Chironna, D. Loconsole, A. Bal, L. Josset, E. Holmes, K. S. George, E. Lasek-Nesselquist, R. S. Sikkema, B. O. Munnink, M. Koopmans, M. Brytting, V. S. Rani, S. Pavani, T. Smura, A. Heim, S. Kurkela, M. Umair, M. Salman, B. Bartolini, M. Rueca, C. Drosten, T. Wolff, O. Silander, D. Eggink, C. Reusken, H. Vennema, A. Park, C. Carrington, N. Sahadeo, M. Carr, G. Gonzalez, S. A. S. Diego, N. V. R. Laboratory, SeqCOVID-Spain, D. C.-. G. Consortium (DCGC), C. D. G. Network (CDGN), D. N. S.-C.-. s. Program, D. o. E. I. Diseases (KDCA), T. d. Oliveira, N. Faria, A. Rambaut, and M. U. G. Kraemer. Tracking the international spread of SARS-CoV-2 lineages B.1.1.7 and B.1.351/501Y-V2 with grinch. Technical Report 6:121, Wellcome Open Research, Sept. 2021. Type: article.

[15] Á. O’Toole, E. Scher, A. Underwood, B. Jackson, V. Hill, J. T. McCrone, R. Colquhoun, C. Ruis, K. Abu-Dahab, B. Taylor, C. Yeats, L. du Plessis, D. Maloney, N. Medd, S. W. Attwood, D. M. Aanensen, E. C. Holmes, O. G. Pybus, and A. Rambaut. Assignment of epidemiological lineages in an emerging pandemic using the pangolin tool. Virus Evolution, 7(2):veab064, Dec. 2021.

[16] F. Pedregosa, G. Varoquaux, A. Gramfort, V. Michel, B. Thirion, O. Grisel, M. Blondel, P. Prettenhofer, R. Weiss, V. Dubourg, J. Vanderplas, A. Passos, D. Cournapeau, M. Brucher, M. Perrot, and É. Duchesnay. Scikit-learn: Machine Learning in Python. Journal of Machine Learning Research, 12(85):2825–2830, 2011.

[17] X. Peng, L. Gralinski, M. T. Ferris, M. B. Frieman, M. J. Thomas, S. Proll, M. J. Korth, J. R. Tisoncik, M. Heise, S. Luo, G. P. Schroth, T. M. Tumpey, C. Li, Y. Kawaoka, R. S. Baric, and M. G. Katze. Integrative Deep Sequencing of the Mouse Lung Transcriptome Reveals Differential Expression of Diverse Classes of Small RNAs in Response to Respiratory Virus Infection. mBio, 2(6):e00198–11, Nov. 2011. Publisher: American Society for Microbiology.

[18] A. Qureshi, N. Thakur, I. Monga, A. Thakur, and M. Kumar. VIRmiRNA: a comprehensive resource for experimentally validated viral miRNAs and their targets. Database, 2014:bau103, Jan. 2014.

[19] A. Rambaut, E. C. Holmes, Á. O’Toole, V. Hill, J. T. McCrone, C. Ruis, L. du Plessis, and O. G. Pybus. A dynamic nomenclature proposal for SARS-CoV-2 lineages to assist genomic epidemiology. Nature Microbiology, 5(11):1403–1407, Nov. 2020. Number: 11 Publisher: Nature Publishing Group.

[20] E. Schubert, J. Sander, M. Ester, H. P. Kriegel, and X. Xu. DBSCAN Revisited, Revisited: Why and How You Should (Still) Use DBSCAN. ACM Transactions on Database Systems, 42(3):19:1–19:21, Apr. 2017.

[21] O. Uǧurel, O. Ata, and D. Balik. An updated analysis of variations in SARS-CoV-2 genome. Turkish Journal of Biology, 44(7):157–167, Jan. 2020.

[22] A. L. Valesano, K. E. Rumfelt, D. E. Dimcheff, C. N. Blair, W. J. Fitzsimmons, J. G. Petrie, E. T. Martin, and A. S. Lauring. Temporal dynamics of SARS-CoV-2 mutation accumulation within and across infected hosts. PLOS Pathogens, 17(4):e1009499, 2021. Publisher: Public Library of Science.

[23] L. Van der Maaten and G. Hinton. Visualizing data using t-SNE. Journal of machine learning research, 9(11), 2008.

[24] A. Varble and B. R. tenOever. Implications of RNA virus-produced miRNAs. RNA Biology, 8(2):190–194, Mar. 2011. Publisher: Taylor & Francis _eprint: https://doi.org/10.4161/rna.8.2.13983.

[25] A. Wu, P. Niu, L. Wang, H. Zhou, X. Zhao, W. Wang, J. Wang, C. Ji, X. Ding, X. Wang, R. Lu, S. Gold, S. Aliyari, S. Zhang, E. Vikram, A. Zou, E. Lenh, J. Chen, F. Ye, N. Han, Y. Peng, H. Guo, G. Wu, T. Jiang, W. Tan, and G. Cheng. Mutations, Recombination and Insertion in the Evolution of 2019-nCoV, Mar. 2020. Pages: 2020.02.29.971101 Section: New Results.

[26] Z. Zhao, B. A. Sokhansanj, C. Malhotra, K. Zheng, and G. L. Rosen. Genetic grouping of SARS-CoV-2 coronavirus sequences using informative subtype markers for pandemic spread visualization. PLoS computational biology, 16(9):e1008269, Sept. 2020.

